# Molecular fate-mapping of serum antibodies reveals the effects of antigenic imprinting on repeated immunization

**DOI:** 10.1101/2022.08.29.505743

**Authors:** Ariën Schiepers, Marije F. L. van ’t Wout, Allison J. Greaney, Trinity Zang, Hiromi Muramatsu, Paulo J. C. Lin, Ying K. Tam, Luka Mesin, Tyler N. Starr, Paul D. Bieniasz, Norbert Pardi, Jesse D. Bloom, Gabriel D. Victora

**Affiliations:** Laboratory of Lymphocyte Dynamics, The Rockefeller University, New York, NY, USA; Basic Sciences Division and Computational Biology Program, Fred Hutchinson Cancer Research Center, Seattle, WA, USA; Laboratory of Retrovirology, The Rockefeller University, New York, NY, USA; Department of Microbiology, Perelman School of Medicine, University of Pennsylvania, Philadelphia, PA, USA; Acuitas Therapeutics, Vancouver, Canada; Howard Hughes Medical Institute, Chevy Chase, MD, USA

## Abstract

The ability of serum antibody to protect against pathogens arises from the interplay of antigen-specific B cell clones of different affinities and fine specificities. These cellular dynamics are ultimately responsible for serum-level phenomena such as antibody imprinting or “Original Antigenic Sin” (OAS), a proposed propensity of the immune system to rely repeatedly on the first cohort of B cells that responded to a stimulus upon exposure to related antigens. Imprinting/OAS is thought to pose a barrier to vaccination against rapidly evolving viruses such as influenza and SARS-CoV-2. Precise measurement of the extent to which imprinting/OAS inhibits the recruitment of new B cell clones by boosting is challenging because cellular and temporal origins cannot readily be assigned to antibodies in circulation. Thus, the extent to which imprinting/OAS impacts the induction of new responses in various settings remains unclear. To address this, we developed a “molecular fate-mapping” approach in which serum antibodies derived from specific cohorts of B cells can be differentially detected. We show that, upon sequential homologous boosting, the serum antibody response strongly favors reuse of the first cohort of B cell clones over the recruitment of new, naÏve-derived B cells. This “primary addiction” decreases as a function of antigenic distance, allowing secondary immunization with divergent influenza virus or SARS-CoV-2 glycoproteins to overcome imprinting/OAS by targeting novel epitopes absent from the priming variant. Our findings have implications for the understanding of imprinting/OAS, and for the design and testing of vaccines aimed at eliciting antibodies to evolving antigens.

## INTRODUCTION

The binding and neutralizing potency of serum antibody is an emergent property of the complex mixture of immunoglobulins secreted over time by B cell clones of various specificities and a range of affinities. The plasma cells that produce such immunoglobulins can arise from multiple parallel pathways. These range from direct differentiation from naïve B cells upon primary infection or immunization to much more complex tracks involving affinity maturation in germinal centers (GCs) and intercalating memory B cell stages, which become progressively more complex over repeated antigenic exposures[1-4]. The contribution of each of these pathways to serum antibody has been difficult to deconvolute: studies of the clonal dynamics of antibody responses rely primarily on molecular analysis of immunoglobulin genes obtained from memory or GC B cells[5-8], and, although direct studies of the clonal composition of antibody in serum have been published[9, 10], current technologies are unable to assign a cellular origin to antibodies of different specificities. Thus, immune phenomena that take place at the serum antibody level are incompletely understood.

One such phenomenon is antigenic imprinting/OAS. OAS was described in the 1950’s as a tendency of individuals exposed to a given strain of influenza to respond with antibodies that reacted more strongly to the first strain of influenza these individuals had met in early childhood than to the exposure strain itself[11, 12]. This was attributed to the immune system’s propensity (which we refer to as “primary addiction”) to repeatedly reuse the first cohort of B cells that respond to an antigen, rather than to recruit new clones from the naïve repertoire[13, 14]. While there is general agreement that imprinting/OAS will impact at least some aspects of immunity to sequentially drifting viruses[15, 16], the full extent to which it impacts future responses to the same or variant antigens is still debated[17, 18]. B cell fate-mapping experiments done in our laboratory[19] show that secondary GCs are composed almost exclusively (> 90%) of naïve, as opposed to memory-derived, B cells. Therefore, at least in mice, primary addiction does not have a pronounced effect at the GC level. Whether this discrepancy is due to a general absence of imprinting or to a divergence between the cellular and serum compartments is not clear.

## RESULTS

Resolving this discrepancy—as well as generally understanding the effect of primary addiction on secondary responses—requires the ability to perform such fate-mapping experiments on serum antibody itself. To achieve this, we adapted the classic fate-mapping approach to enable detection of the cellular and temporal origin of antibodies in serum, an approach we call “molecular fate-mapping.” We engineered mice in which the C-terminus of the immunoglobulin kappa (Igκ) light chain gene (*Igk*) is extended to encode for a LoxP-flanked FLAG-tag, followed by a downstream Strep-tag (**Fig. 1a and Supplementary Fig. 1**). B cells bearing this “Κ-tag” allele produce immunoglobulins that are FLAG-tagged unless exposed to Cre recombinase, upon which they permanently switch the FLAG-tag for a Strep-tag. Cre-mediated recombination thus fate-maps the antibodies these B cells and their plasma cell descendants express on their surfaces and/or secrete into serum. This allows for differential detection of pre- and post-fate-mapping Igκ^+^ antibody using secondary reagents specific for each tag.

**Figure 1.**
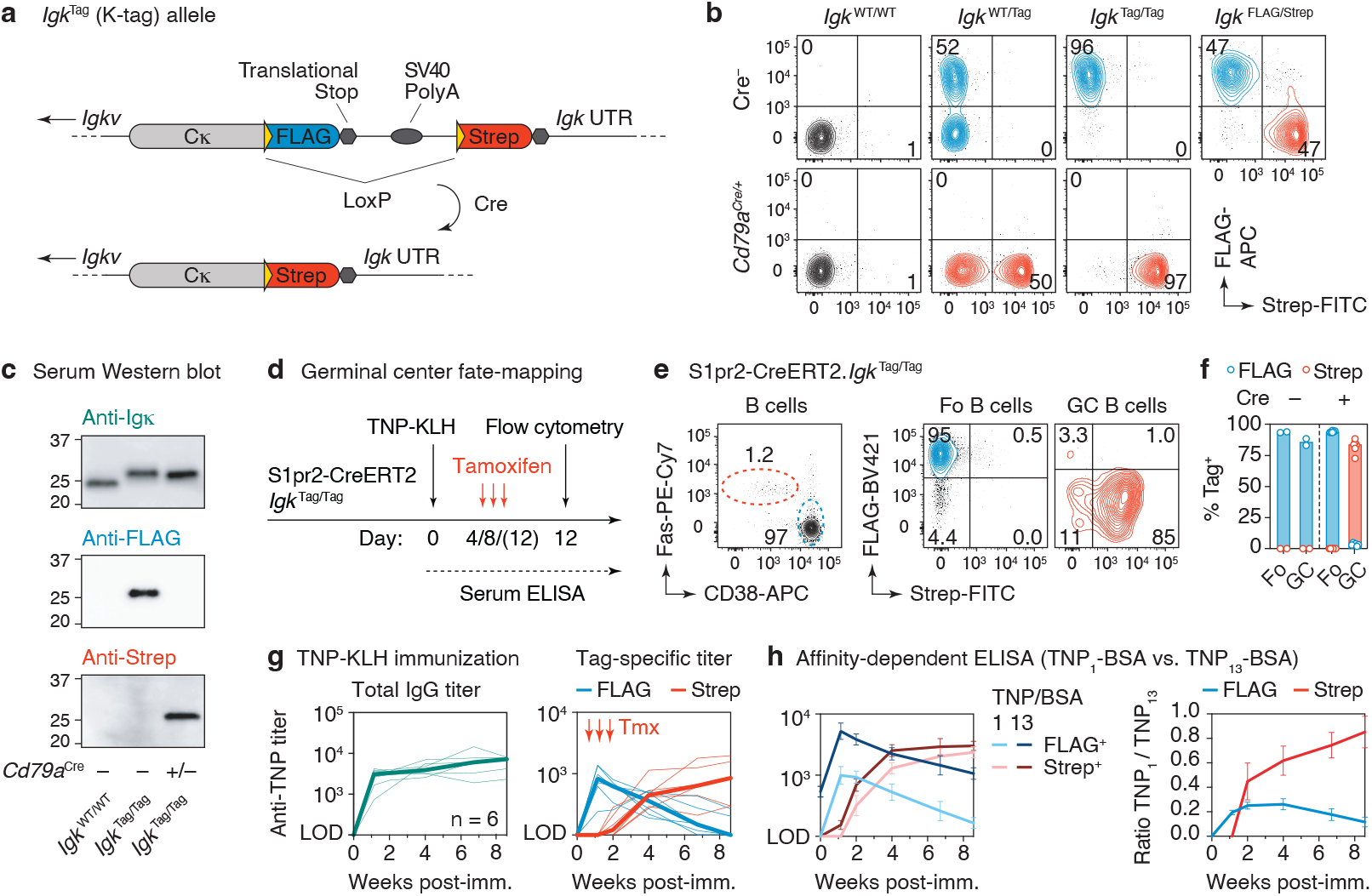
The Κ-tag system for molecular fate-mapping of serum antibodies. **(a)** Schematic representation of the *Igk*^Tag^ allele prior to and following cre-mediated recombination. **(b)** Flow cytometry of blood B cells showing expression of FLAG- and Strep-tagged B cell receptors on mice of the indicated genotype. **(c)** Western blotting of serum obtained from mice of the indicated genotype, stained for Igκ light chain or FLAG/Strep-tags. **(d)** Schematic representation of the immunization strategy used to fate-map GC B cells and their antibody output. **(e)** Flow cytometry of popliteal lymph node 12 days after footpad immunization with TNP-KLH in alum adjuvant. Left panel shows B cells (B220^+^, CD4^-^ CD8^-^ CD138^-^) stained for GC (FAS^+^ CD38^−^) and follicular (Fo) B cell (FAS^−^ CD38^+^) markers. **(f)** Quantification of data in (e). Each dot represents an individual lymph node, bars represent the median. **(g)** Anti-TNP total IgG and tag-specific endpoint titers as determined by TNP_4_ -BSA ELISA in mice immunized i.p. with TNP-KLH in alhydrogel adjuvant. Thin lines represent individual mice, thick lines link the medians of the log transformed titer values at each time point. **(g)** Relative affinity of anti-TNP FLAG^+^ and Strep^+^ antibodies as estimated by ELISA using TNP_1_-BSA or TNP_13_-BSA as capture reagents. The means of the log transformed titer values are shown with error bars representing SEM. The ratio between anti-TNP_1_-BSA and anti-TNP_13_-BSA titers was calculated per sample, shown in the right panel.

To verify the functionality of the Κ-tag allele, we first determined that B cells in *Igk*^Tag^ mice expressed tagged B cell receptors on their surface. Following the rules of allelic exclusion[20], roughly 50% and 95% of B cells from heterozygous *Igk*^WT/Tag^ (WT, wild-type) or homozygous *Igk*^Tag/Tag^ mice were FLAG^+^ (as expected, ∼5-10% of B cells in homozygous mice carried an Igλ light chain[20]; **Supplementary Fig. 2a**). Κ-tag mice in which all B cells constitutively expressed Cre recombinase (*Igk*^Tag/Tag^.*Cd79a*^Cre/+^) replaced FLAG-with Strep-tag in virtually all B cells (**Fig. 1b**). Importantly, Κ-tag mice appropriately secreted FLAG- or Strep-tagged antibodies into serum in the absence or presence of Cre recombinase, respectively (**Fig. 1c**), without affecting steady state serum antibody levels (**Supplementary Fig. 2b**). Generation and maturation of Κ-tag B cells was unimpaired, as indicated by the equal proportion of tagged and untagged circulating B cells in *Igk*^WT/Tag^ mice (**Fig. 1b**). The same was true of circulating B cells and bone marrow plasma cells expressing FLAG- and Strep-tags in *Igk*^FLAG/Strep^ mice, in which one of the two Κ-tag alleles was pre-recombined by Cre expression in the germline (**Fig. 1b** and **Supplementary Fig. 2c**).

To ensure that FLAG^+^ and Strep^+^ B cells were equally competitive throughout the course of B cell activation, affinity maturation, and plasma cell differentiation, we primed and boosted *Igk*^FLAG/Strep^ mice with the model antigen 2,4,6-trinitrophenyl-keyhole limpet hemocyanin (TNP-KLH) in alum adjuvant and followed the serum titers of antibodies bearing each tag over time. To enable direct comparison of titers of anti-TNP antibodies bearing each tag, we diluted secondary (anti-FLAG or anti-Strep) antibodies to achieve similar detection levels on standard curves generated using recombinant FLAG- or Strep-tagged monoclonal antibodies (**Supplementary Fig. 2d**). Endpoint ELISA titers normalized using these curves showed a similar range of anti-TNP reactivity between FLAG^+^ and Strep^+^ fractions (**Supplementary Fig. 2e**), indicative of equal competitiveness of the differently tagged B cells.

Following the serum antibody produced by B cells engaged at different stages of the immune response requires temporally restricted fate-mapping of activated B cell clones. To enable this, we crossed *Igk*^Tag^ mice to the GC-specific, tamoxifen-inducible S1pr2-CreERT2 BAC-transgenic allele[21] (to generate S1pr2-*Igk*^Tag^ mice). Tamoxifen treatment of mice on days 4 and 8 after TNP-KLH immunization led to efficient recombination (96.1% +/-0.50 SEM ((Strep^+^/Tag^+^) x 100)) of the Κ-tag allele in GC B cells but not in non-GC B cells in the same lymph node at 12 days post-immunization (d.p.i.) (**Fig. 1d-f**). Again, tagged B cells were found at similar proportions to untagged B cells in heterozygous S1pr2-*Igk*^WT/Tag^ mice (on average 41% +/-SD 9.0 Tag^+^), indicating that expression of the tag does not impair B cell competitiveness in the GC (**Supplementary Fig. 2f, g**). GC B cells in Cre^−^ animals (**Fig. 1f**) or S1pr2-*Igk*^WT/Tag^ mice not treated with tamoxifen remained FLAG^+^, with only minimal spontaneous recombination (1.3% ± 1.0 SD Strep^+^ at day 12 d.p.i.) in the latter (**Supplementary Fig. 2g**).

Total anti-TNP IgG antibodies in S1pr2-*Igk*^Tag^ mice immunized intraperitoneally (i.p.) with TNP-KLH in alhydrogel and treated with tamoxifen on days 4, 8 and 12 were first detected in serum at 8 d.p.i. and increased progressively through 60 d.p.i. (**Fig. 1g**). Deconvolution of GC-derived (Strep^+^) and non-GC derived (FLAG^+^) antibodies showed that an initial wave of extrafollicular FLAG^+^ antibodies that peaked at 8 d.p.i. was progressively replaced by GC-derived Strep^+^ antibodies that were first detected in serum at 14 d.p.i. (**Fig. 1g**). FLAG^+^ anti-TNP antibodies regressed to near baseline levels between 47-60 d.p.i., as expected from their extrafollicular origin. An affinity-dependent anti-TNP ELISA showed detectable affinity maturation only in the GC-derived Strep^+^ antibody fraction (**Fig. 1h**), confirming the efficient fate-mapping of GC-derived antibody. Background signal in control animals not given tamoxifen remained below the limit of detection (LOD) throughout the primary response (**Supplementary Fig. 2h**). Thus, the S1pr2-*Igk*^Tag^ mouse model enables us to discern antibodies derived from the first wave of B cells that entered a GC reaction in response to immunization. Our data also show that extrafollicular responses are of relatively short duration in these settings, and that virtually all antibody detectable after the first few weeks of immunization, and especially antibody with high affinity, is derived from plasma cells of GC origin.

We next leveraged the ability to trigger Cre-mediated recombination of the Κ-tag allele in a time-resolved manner to mark serum antibodies produced by B cells that formed GCs in response to primary immunization (the “primary cohort”) and follow their kinetics upon boosting. In this setting, primary cohort-derived antibodies are Strep^+^, whereas new antibody originating from naïve B cells (as well as any non-GC-derived primary memory B cells) engaged by the boost carry a FLAG-tag. This approach allows us to measure primary addiction at “zero-antigenic distance”— i.e., when priming and boosting use the exact same antigen—thus avoiding confounding by a potential negative effect of antigenic distance on the degree of imprinting. We first primed S1pr2-*Igk*^Tag^ mice i.p. with alum-adjuvanted TNP-KLH and administered tamoxifen at 4, 8 and 12 d.p.i. to fate-map primary cohort GC B cells and their memory and plasma cell progeny (**Fig. 2a**).

**Figure 2.**
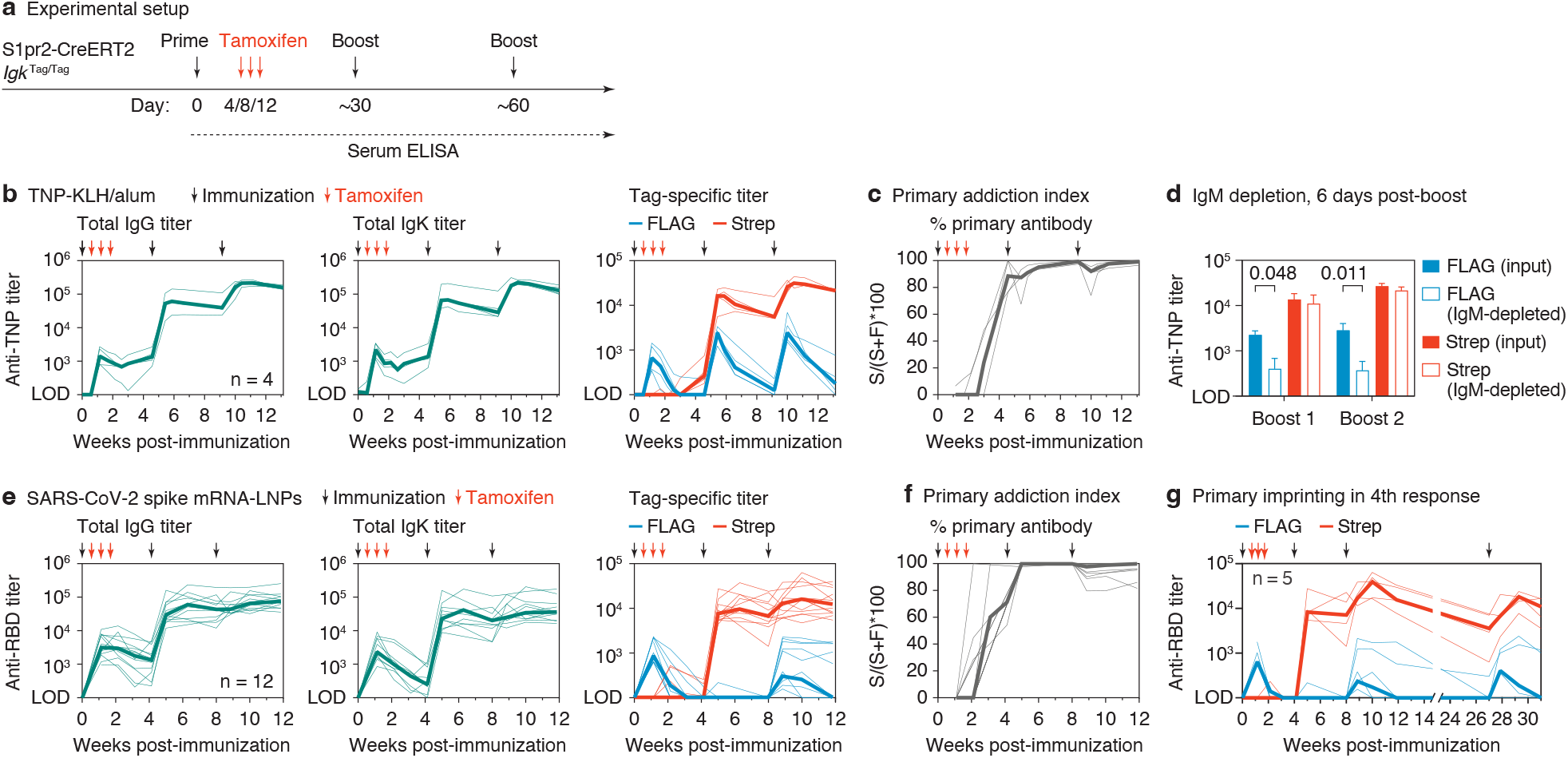
Primary addiction upon homologous boosting. **(a)** Schematic representation of general immunization strategy used to measure primary addiction. S1pr2-*Igk*^Tag/Tag^ mice were immunized on the days indicated by black arrows with TNP-KLH in alum (b-d) or WH1 mRNA-LNP (e-g) and treated with tamoxifen at 4, 8 and 12 d.p.i. as indicated by the red arrows. **(b)** Anti-TNP serum IgG (left panel), Igκ (middle panel) and tag-specific titers (right panel) as measured using TNP_4_ -BSA by ELISA. Results are from 4 mice from 2 independent experiments. Thin lines represent individual mice, thick lines link the medians of the log transformed titer values at each time point. **(c)** The percentage of the TNP-titer derived from the primary cohort of B cells (the “primary addiction index”) was calculated by dividing the Strep^+^ titer of each individual sample by its total titer (Strep^+^ + FLAG^+^), multiplied by 100 (S/(S+F)*100). **(d)** Tag-specific anti-TNP titers before and after IgM depletion for samples collected 6 days after the first and second boost with TNP-KLH. Bars represent the means of log transformed titers and the error bars are standard errors of the mean. P-values are for paired T-test, only statistically significant (p < 0.05) values are shown. **(e)** Anti-WH1 RBD IgG (left panel), Igκ (middle panel), and tag-specific titers (right panel) as measured by ELISA. Results are from 12 mice from 3 independent experiments. **(f)** Primary addiction index calculated as in (c). **(g)** Anti-WH1 RBD response in mice receiving a 4^th^ dose of mRNA-LNP at 133 days after the previous dose, for one of the cohorts shown in (e). Two of five mice were not sampled at day 0.

Boosting with the same antigen one and two months after the primary immunization resulted in the expected increases in recall TNP titers (**Fig. 2b**). Deconvolution of these responses using tag-specific ELISA revealed that both secondary and tertiary titers were strongly dominated by fate-mapped (Strep^+^) antibodies derived from primary cohort B cells and, whereas FLAG^+^ titers also increased after each boost, these peaked at much lower levels and decayed markedly with time (**Fig. 2b**). A “primary addiction index” computed by dividing Strep^+^ by total Strep^+^ + FLAG^+^ titers (S/(S+F)*100) showed that almost all detectable recall antibody (mean 95% and 97% of serum reactivity at 14 days after the first and second boosts, respectively) was derived from the B cell cohort engaged in the primary GC response (**Fig. 2c**), despite secondary GCs being composed of new (FLAG^+^) B cells ([19] and **Supplementary Fig. 3a**). Depletion of IgM from post-boost serum samples (**Supplementary Fig. 3b**) resulted in a sharp reduction in FLAG^+^ but not Strep^+^ recall TNP titers, supporting the notion that naïve-derived B cells engaged by recall generate primarily an extrafollicular-like (IgM-dominated) B cell response (**Fig. 2d**).

To extend these findings to a disease-relevant setting, we immunized and boosted mice as in **Fig. 2a** but using a lipid nanoparticle (LNP)-formulated nucleoside-modified mRNA vaccine encoding the prefusion-stabilized (2P) form of the SARS-CoV-2 Wuhan-Hu-1 (WH1) spike (S) protein, similar to available SARS-CoV-2 mRNA vaccines[22]. Secondary and tertiary anti-S protein receptor binding domain (RBD) antibodies were again almost entirely derived from primary cohort (Strep^+^) B cells (**Fig. 2e,f**). As with GC B cells (**Supplementary Fig. 2g**), low-level spontaneous recombination to Strep^+^ antibody was detected in recall responses in control mice not given tamoxifen. This resulted in a slight underestimation of FLAG^+^ antibody titers (median 3.9 and 3.6 two weeks after second and third immunizations respectively; **Supplementary Fig. 3c**). Although primary addiction was more pronounced than for the SARS-CoV-2 RBD than for TNP-KLH after the first boost (no new (FLAG^+^) antibody was detected at this time point), 5/12 mice developed low but stable titers of FLAG^+^ anti-RBD antibody after the third dose. This bimodality was independent of the experimental cohort and of whether boosting was done ipsilaterally or contralaterally to the site of the primary dose (**Supplementary Fig. 3d**) and is therefore likely ascribable to stochastic variability inherent to highly oligoclonal recall responses[19]. Primary addiction was long-lasting, as even a fourth immunization of a subset of mice (>133 days after the previous boost) was dominated by Strep-tagged antibodies (**Fig. 2g and Supplementary Fig. 3e,f**). Together, our experiments reveal a high degree of primary addiction in recall responses induced by homologous boosting across different immunization models. We conclude that imprinting/OAS can be remarkably strong when measured at zero antigenic distance.

To define how primary addiction responds to the antigenic distance between priming ant boosting antigens, we employed historical series of drifted influenza virus hemagglutinin (HA) variants as models. We first used an influenza infection/immunization model based on the two strains for which the OAS phenomenon was originally described—A/Puerto Rico/8/1934 (PR8) and A/Fort Monmouth/1/1947 (FM1)[11, 12]—whose HAs share 90% identity at the amino acid level (Fig. 3b). Following the schematic in **Fig. 3a**, we infected S1pr2-*Igk*^Tag^ mice with mouse-adapted influenza PR8 virus, fate-mapped the primary cohort of GC B cells with 4 doses of tamoxifen given between 4 and 16 days post-infection, then followed the evolution of primary (Strep^+^) and new (FLAG^+^) antibody fractions over time. As with immunization, the primary response to HA_PR8_ was characterized by high extrafollicular (FLAG^+^) titers that peaked between 8 and 16 days post-infection and were subsequently replaced by GC-derived (Strep^+^) titers (**Fig. 3c**). Homologous boosting with recombinant HA_PR8_ protein subcutaneously at 3 and 4 months post-infection resulted in a 1-log increase in Strep^+^ HA_PR8_ binding titers after the first boost and a less pronounced increase after the second boost. As with protein immunization, the contribution of non-primary (FLAG^+^) antibodies to total titers was minor: even though it increased progressively between the first and second boosts, its peak median value was roughly 10% of HA reactivity (**Fig. 3c**). Heterologous boosting with HA_FM1_ led to only slight back-boosting of primary Strep^+^ HA_PR8_ titers and had virtually no effect on FLAG^+^ HA_PR8_ reactivity (**Fig. 3d**), indicative of substantial antigenic distance between these variants. Accordingly, crossreactive primary titers towards HA_FM1_ were completely absent from the primary extrafollicular response to PR8 infection and began to emerge only at approximately 4 weeks in the Strep^+^ antibody fraction (**Fig. 3d**), likely as a side-effect of affinity maturation towards HA_PR8_. Heterologous boosting not only increased these crossreactive (Strep^+^) titers by close to 1 log, but, importantly, also induced substantial responses from non-OAS boost-elicited B cell clones, in that roughly half of all serum reactivity to HA_FM1_ was derived from the FLAG^+^ fraction after the second boost (**Fig. 3d**). Thus, heterologous boosting can partly circumvent primary addiction, leading to the expansion of variant-specific B cell clones not involved in the primary response.

**Figure 3.**
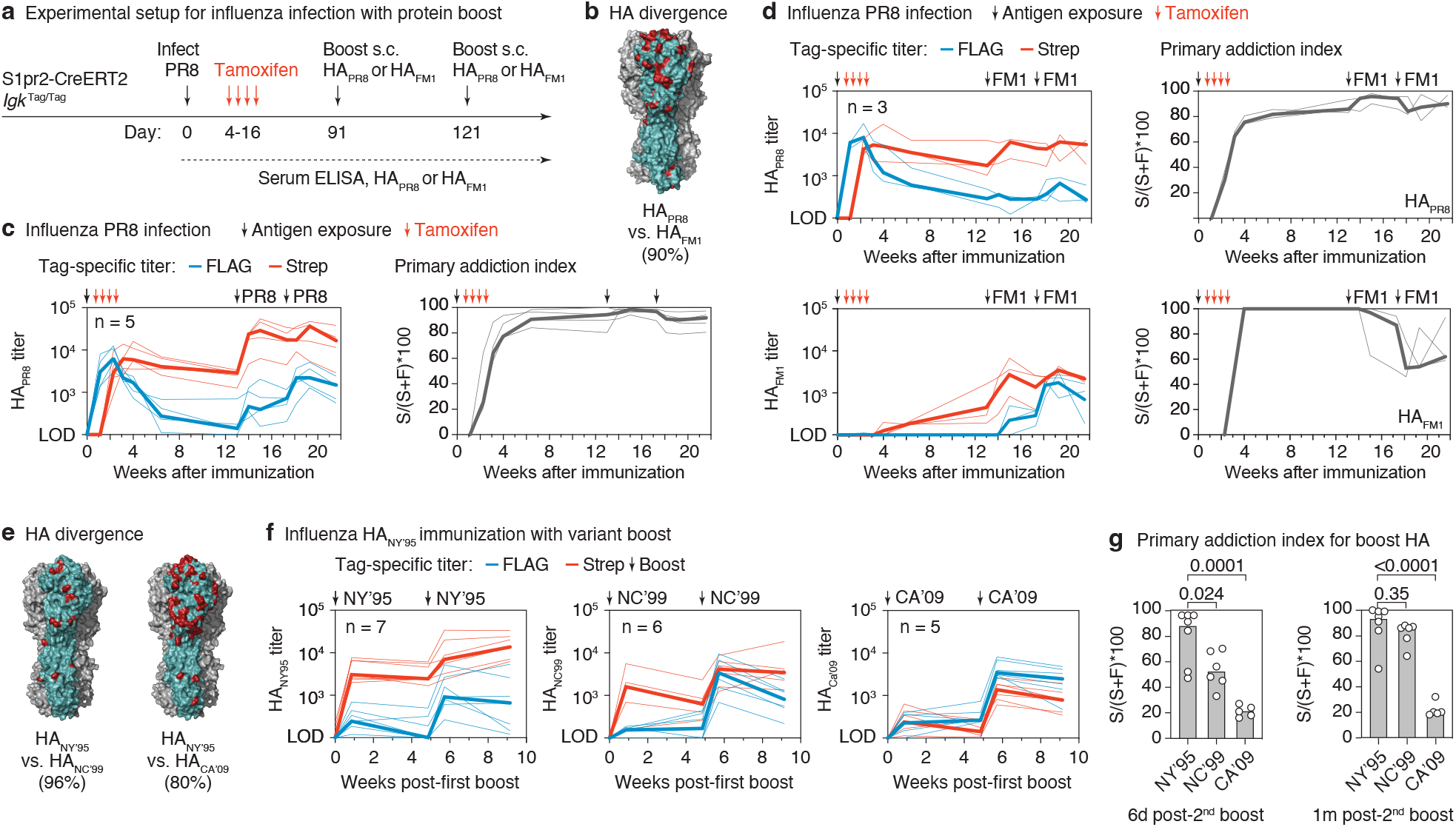
Primary addiction decreases with antigenic distance. **(a)** Schematic representation of influenza infection and HA boosting strategy. S1pr2-*Igk*^Tag/Tag^ mice were infected intranasally with PR8 influenza and boosted subcutaneously (s.c.) with HA_PR8_ or HA_FM1_ in alhydrogel at the indicated time points. **(b)** Rendering of the HA_PR8_ trimer structure (PDB: 1RU7) with one monomer highlighted in teal and the amino acids that diverge between HA_PR8_ and HA_FM1_ in red. Percent aa identity is given in parentheses. **(c)** Anti-HA_PR8_ FLAG^+^ and Strep^+^ titers in S1pr2-*Igk*^Tag/Tag^ mice homologously boosted with HA_PR8_ (left) and quantification of the primary addiction index score (right). **(d)** Anti-HA_PR8_ (top) and anti-HA_FM1_ (bottom) ELISA reactivity in mice heterologously boosted with HA_FM1_. Tag-specific titers are shown in the left panels, and quantification of the primary addiction index score is show on the right. **(e)** Divergence between HA_NC’95_ and HA_NC’99_ or HA_CA’09_, colored as in (b). Modeled on the structure of HA_CA’09_ (PDB: 3LZG). **(f)** Anti-HA tag-specific titers in mice primed with HA_NY’95_ protein, shown after the first boost with HA_NY’95_ (homologous) or with variants HA_NC’99_ or HA_CA’09_ (heterologous), as outlined in Supplementary Fig. 4a. Antibody reactivity against the boosting antigen is shown. The full time-course and reactivities against all three HAs as measured by ELISA for the top serum dilution are shown in Supplementary Fig. 4b. **(g)** Primary addiction index for the boosting HA, 6 days (left) and 1 month (right) after the second boost (third dose). P-values are for Student’s T test comparing the primary addiction index of the homologous to each heterologous boost. Thin lines represent individual mice, thick lines link the medians of the log transformed titer values at each time point.

To verify this notion over a wider range of antigenic distances, we immunized mice i.p. with recombinant H1 from strain A/New York/614/1995 (NY/95) in alhydrogel adjuvant, then boosted these mice twice, either homologously with NY/95 or heterologously with H1s from strains A/New Caledonia/20/1999 (NC/99; a slightly drifted strain with 96% amino acid identity to NY/95) or pandemic A/California/07/2009 (CA/09; an “antigenic shift” strain, with 80% amino acid identity, **Fig. 3e and Supplementary Fig. 4a**). Generally, primary addiction was weaker and more variable in this setting even to the homologous response, possibly due to the overall weak primary response elicited by immunization with recombinant HA_PR8_ protein (**Supplementary Fig. 4b)**. Nevertheless, we observed a progressive decrease in primary addiction as the antigenic distance between the primary and boost antigens increased, so that up to 80% of total serum responses to CA/09 were FLAG-tagged (corresponding to 20% primary addiction) upon boosting with this variant (**Fig. 3f,g**). We conclude that increases in antigenic distance between priming and boosting antigens can decrease imprinting/OAS, enabling the generation of new, variant-specific responses.

A clinically important setting in which to measure the ability of a drifted variant to overcome imprinting/OAS is the response to Omicron strains of SARS-CoV-2 in individuals previously exposed to WH1 SARS-CoV-2 antigens. We thus used the K-tag system to estimate the degree to which boosting with nucleoside-modified mRNA-LNP encoding the S protein from the Omicron BA.1 strain of SARS-CoV-2 was capable of overcoming imprinting/OAS generated by priming with wild-type S-encoding mRNA-LNP (WH1 and BA.1 strains have 98% and 92% amino acid identity in the full S protein and RBD domains, respectively). We primed S1pr2-*Igk*^Tag^ mice with WH1 mRNA-LNP in the right leg, then boosted these mice one and two months later with either BA.1 or WH1 mRNA-LNP distally in the left leg (**Fig. 4a**). Boosting induced similar total IgG responses to WH1 and BA.1 RBDs in both groups (**Fig. 4b**). Tag-specific ELISAS showed that responses to the WH1 RBD were indistinguishable between homologously and heterologously boosted animals, in that primary (Strep^+^) antibodies were strongly dominant in both settings, with no substantial FLAG^+^ response after the initial extrafollicular response (**Fig. 4c,d**). Notably, whereas little to no Strep^+^ antibodies to the BA.1 RBD were observed after primary immunization, recall Strep^+^ reactivity to the BA.1 RBD was equally strong regardless of which variant was used for boosting (**Fig. 4c,d**). This observation agrees with previous reports regarding the evolution of crossreactivity to other strains as a consequence of affinity maturation towards WH1 RBD mRNA vaccination in humans^24^. Importantly, however, heterologous boosting resulted in a pronounced increase in BA.1 RBD titers generated by newly recruited (FLAG^+^) clones not crossreactive with the WH1 strain, a reactivity otherwise absent from mice boosted homologously (**Fig. 4c,d**). At their peak (2 weeks post 2^nd^ boost), Strep^+^ antibodies accounted for an average of 73% (± 19% SD) of total anti-BA.1 reactivity across heterologously boosted mice (**Fig. 4d**). The induction of new (FLAG^+^) antibodies upon double BA.1 boost was even more pronounced when assaying for reactivity against the full-length WH1 and BA.1 S proteins (**Fig. 4e**). Thus, the key difference between boosting homologously and heterologously is that only the latter is capable of eliciting a robust response to the drifted strain by naïve B cells.

**Figure 4.**
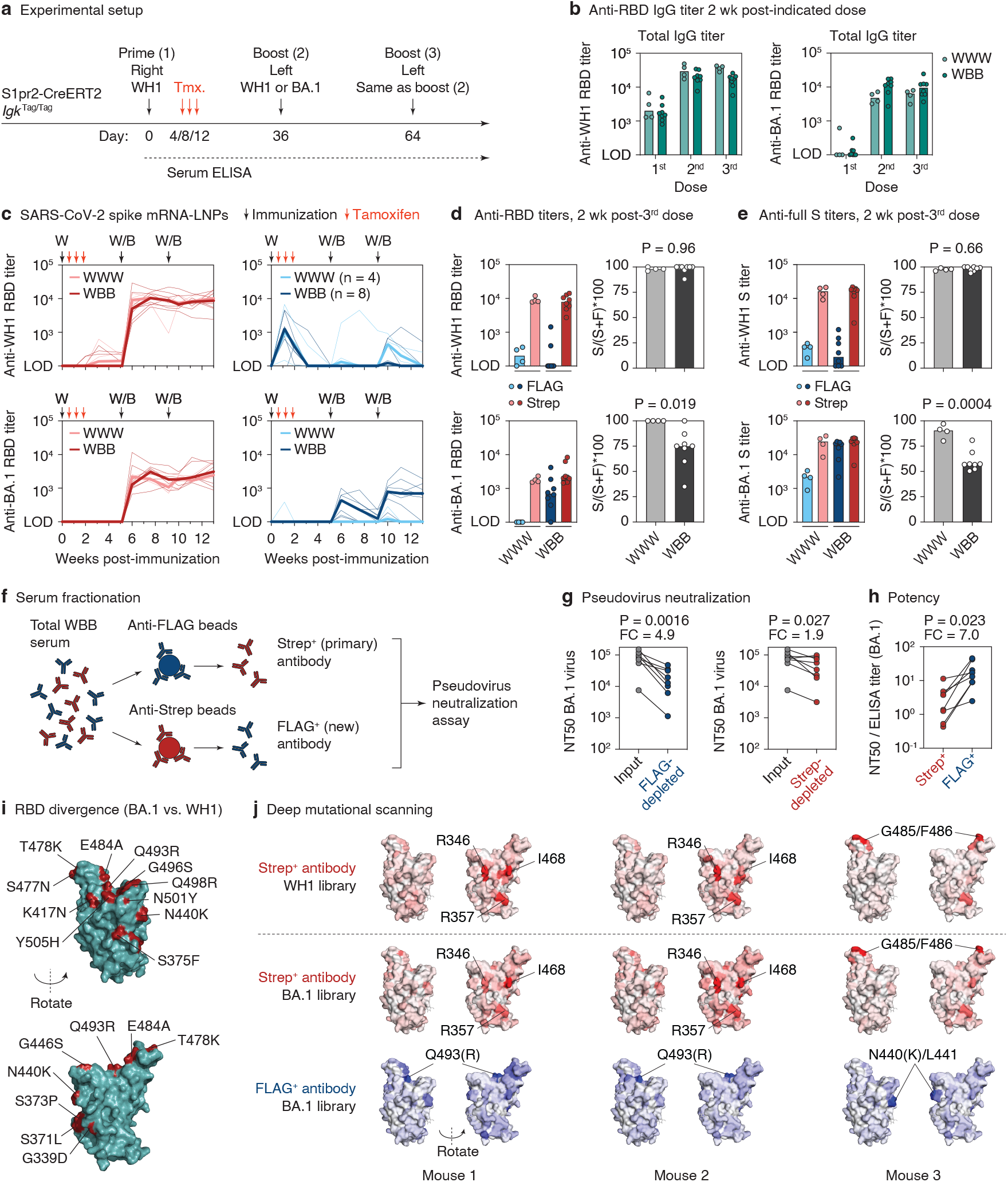
Subversion of primary addiction by heterologous SARS-CoV-2 boosting. **(a)** Schematic representation of homologous and heterologous immunization strategies used to measure primary addiction. As in Fig. 2e, with all mice boosted contralaterally. **(b)** Anti-WH1 and BA.1 RBD IgG titers after the first, second and third doses in homologously (WWW) or heterologously (WBB) immunized S1pr2-*Igk*^Tag/Tag^ mice. Results are from 2 independent experiments. **(c)** Evolution of anti-WH1 and BA.1 RBD tag-specific titers over time. Thin lines represent individual mice, thick lines link the medians of the log transformed titer values at each time point. **(d)** Comparison of FLAG and Strep anti-RBD titers shown in (c) at 2 weeks after the third immunization (left) along with primary addiction index (right). P-values are for Student’s T test. **(e)** Anti-full S tag-specific titers for the same samples, as in (c). **(f)** Schematic representation of the antibody fractionation protocol. **(g)** Thin lines represent individual mice, thick lines link the medians of the log transformed titer values at each time point. 50% pseudovirus neutralization (NT50) titers for WBB mice at the time point shown in (d). Full curves shown in Supplementary Fig. 5b. Post-depletion NT50s were normalized to input serum based on the BA.1 RBD ELISA titer in Supplementary Fig. 5a. P-values are for one-tailed paired T test. FC, fold-change. **(h)** Potency of anti-RBD BA.1 antibody Strep^+^ (FLAG-depleted) and FLAG^+^ (Strep-depleted) fractions, unnormalized NT50 values from the graphs in Supplementary Fig. 5b were divided by the ELISA titer of their respective depletion fraction shown in Supplementary Fig. 5a. P-values are for two-tailed paired T test. FC, fold-change. **(i)** Rendering of the WH1 RBD structure (PDB: 6MOJ) with amino acid changes in BA.1 highlighted in red. **(j)** Deep mutational scanning results of serum samples obtained from 3 heterologously immunized mice at 2 weeks after the third immunization (d). Antibody binding sites on the RBD are shaded according to the escape fraction. The positions most highly targeted by each serum fraction are indicated.

Because viral escape mutations are likely to accumulate in positions important for antibody-mediated neutralization, we speculated that BA.1-specific B cell clones elicited by heterologous boosting may preferentially target escape epitopes, thus producing antibodies that are more potent neutralizers of the drifted strain. To test this, we fractionated serum samples from 14 days after the third immunization into FLAG-depleted (Strep^+^, *primary*) and Strep-depleted (FLAG^+^, *new*) antibody preparations (**Fig. 4f** and **Supplementary Fig. 5A**) and measured their potency against WH1 and BA.1 strains in a pseudotyped virus neutralization assay[24]. As expected, the effect of FLAG^+^ antibody depletion on the neutralization of WH1 virus by either cohort was minimal, whereas Strep^+^ antibody depletion resulted in large decreases in neutralization (1.7 vs 8.5-fold decreases, respectively; **Supplementary Fig. 5b,c**). Thus, BA.1 mRNA-LNP were able to “back-boost” WH1-induced primary memory B cells that were potent WH1 neutralizers. By contrast, removing new (FLAG^+^) antibody from BA.1-boosted sera resulted in a greater reduction of BA.1 neutralization than removing primary (Strep^+^) antibody from these same samples (4.9 vs 1.9-fold decrease; **Fig. 4g**). To estimate the potency per unit of specific antibody, we divided the 50% neutralization titer (NT50) derived from the BA.1 pseudovirus assay by the endpoint binding titer obtained by BA.1 RBD-specific ELISA. When normalized to reactivity, new (FLAG^+^) antibody was on average 7.0-fold more powerful at neutralizing BA.1 than the Strep^+^ antibody produced by B cell clones elicited by WH1 immunization (**Fig. 4h)**.

To understand this difference at the epitope level, we carried out deep mutational scanning[25, 26] to define BA.1 mutants that escaped binding to FLAG^+^ and Strep^+^ antibodies from serum samples of heterologously boosted mice displaying high Strep^+^ and FLAG^+^ ELISA titers (**Fig. 4i and Supplementary Fig. 6)**. This approach allowed us to separately determine the epitopes targeted by primary and new antibody in the same mouse. We measured the antibody escape patterns of four mice against BA.1 (both Strep^+^ and FLAG^+^ antibodies) and WH1 RBD (Strep^+^ antibodies only), three of which showed interpretable dominance patterns (**Supplementary Fig. 7**). In these three mice, there was clear segregation of the epitopes targeted by primary Strep^+^ and new FLAG^+^ antibodies (**Fig. 4j** and **Supplementary Fig. 7)**. In two mice, Strep^+^ antibodies targeted residues of the “class 3” epitope located on the outer face of the RBD (R346, R357, I468), which, as expected, were conserved between WH1 and BA.1 (**Fig. 4i,j)**. By contrast, FLAG^+^ antibodies in both mice were focused on the BA.1-specific residue R493 (Q493 in WH1), located on the top of the RBD in the “class 2” region of the ACE2-binding surface. The third mouse showed a similar pattern of segregation between primary and new antibodies, but targeted different epitopes. Whereas Strep^+^ antibodies were heavily focused on the conserved class 1/2 epitope that includes G485/F486 (on the “top” of the RBD at the ACE2 interface), FLAG^+^ antibodies, targeted primarily an epitope that includes the BA.1-specific K440 residue (N440 in WH1) on the side face of the RBD distal to G485/F486 (class 3). Notably, both N440K and Q493R have been reported to lead to escape from neutralization by various monoclonal antibodies[27-29]. Thus, new antibodies elicited by heterologous immunization were found to preferentially target epitopes that contain BA.1-specific escape mutations and that do not overlap with epitopes bound by crossreactive primary antibodies.

## DISCUSSION

Together, our findings using the Κ-tag system make two major points: first, the ability to measure imprinting/OAS at zero antigenic distance revealed that, in most repeated immunization regimens, the large majority of serum antibody is expected to derive from the primary cohort of responder B cells, even after multiple rounds of boosting. Given our previous observation that secondary GCs in mice consist almost exclusively of naïve-derived B cell clones[19], these findings indicate a functional divide between cellular and serum responses, where only the latter are dominated by the effects of imprinting/OAS. A potential explanation for this divergence is that the naïve B cells that contribute to secondary GCs are not of sufficient affinity either to exit the GC as plasma cells or to secrete antibody that is detectable by direct ELISA. Low antigen binding among naïve-derived secondary GC B cells has been reported previously by others[30], which is consistent with the latter hypothesis.

Second, our experiments using the influenza virus and SARS-CoV-2 models show that imprinting/OAS is sensitive to the antigenic distance between the priming and boosting strains. Of note, we found OAS in our infection/immunization model to be relatively weak between the PR8 and FM1 influenza virus strains with which this phenomenon was described[11, 12], possibly accounting for the ability of FM1 vaccination to overcome PR8 imprinting in humans[31]. Because antigenic drift for both influenza virus and SARS-CoV-2 is primarily driven by immune escape, loosening of primary addiction should in principle focus newly recruited B cell clones on drifted neutralizing epitopes, a view supported by the results of our neutralization and epitope mapping experiments. Moreover, antibodies that escaped imprinting tended to focus on novel residues located within epitopes that were not dominant targets of the crossreactive first-cohort response. This pattern suggests antibody-mediated epitope masking—whereby serum antibody either present prior to boosting or produced acutely by boosted memory B cells competes with naïve B cells for binding a specific epitope—as a potential mechanism for imprinting/OAS. Similar effects have been observed previously by infusion of monoclonal antibodies prior to induction of an immune response in mice and humans and are predicted to affect the fine-specificity of recall responses[32-34]. These findings also suggest a teleological explanation for why it might be advantageous for memory B cells to avoid re-entering secondary GCs[19, 35], since competition by memory B cells could inhibit the ability of naïve cells to generate antibodies tailored to novel epitopes in viral escape variants. In such a framework, the primary role of secondary GCs would be to circumvent the worst effects of OAS/imprinting.

Our observation that mice boosted homologously with WH1 mRNA had only slightly (and not significantly) lower whole-IgG titers against RBDs of both strains than mice boosted with BA.1 mRNA is in agreement with data from non-human primates in showing that neutralizing antibodies to both strains are similarly induced by homologous and heterologous boosting[36]. The same study found similar expansion of cross-reactive memory B cells in both groups, as also reported for Omicron breakthrough infections in WH1-vaccinated humans[37, 38]. However, deconvolution of antibody responses into primary and boost-elicited clones using the K-tag system revealed that heterologous boosting was uniquely capable of recruiting new, BA.1-specific B cell clones with greater neutralizing potency towards the variant strain. Importantly, a second dose of BA.1 was required to amplify this new antibody reactivity, most likely because it directed memory B cells generated by the first BA.1 boost into the plasma cell compartment. Barring major differences in the logic of imprinting/OAS between humans and mice, our findings suggest that Omicron-specific boosters will induce B cell clones that are superior neutralizers of Omicron strains, although their full effectiveness may be best measured after a second heterologous booster dose. Our findings therefore contribute to the ongoing debate on the value of Omicron-specific boosters for the broadening of protection to SARS-CoV-2.

## Supporting information

DMS results

## Acknowledgments

We would like to thank the Rockefeller University Transgenics and Gene Targeting facilities for generating the K-tag mouse strain and Comparative Biosciences Center for mouse housing. We thank P. Wilson (Weill Cornell Medicine) for recombinant SARS-CoV-2 WH1 Spike and RBD protein, J.T. Jacobsen (Rockefeller University) for technical assistance, and all Rockefeller University staff for their continuous support. This study was funded by NIH/NIAID grants R01AI119006 and R01AI139117 to G.D.V., P01AI165075 to P.D.B., R01AI146101 and R01AI153064 to N.P., and R01AI141707 to J.D.B. Work in the Victora laboratory is additionally supported by NIH grant DP1AI144248 (Pioneer award) and the Robertson Foundation. A.S. was supported by a Boehringer-Ingelheim Fonds PhD fellowship. P.D.B. and J.D.B. are HHMI investigators. G.D.V. is a Burroughs-Wellcome Investigator in the Pathogenesis of Infectious Disease, a Pew-Stewart Scholar, and a MacArthur Fellow.

## Author Contributions

A.S. and M.F.L.v.’t.W. performed all experimental work, with essential input from L.M.. A.J.G. performed and interpreted the deep mutational scanning experiments under supervision of T.N.S. and J.D.B.. N.P. and H.M. designed and produced WH1 and BA.1 S-encoding mRNAs, P.J.C.L. and Y.K.T. formulated mRNAs into LNP. T.Z. carried out pseudovirus neutralization assays under supervision of P.D.B.. A.S. and G.D.V. designed the K-tag allele and all of the experiments in the study and wrote the manuscript with input from all authors.

## Data Availability

The raw Illumina reads of the 16-nucleotide variant barcodes from the deep mutational scanning experiments are available on the NCBI SRA under BioProject PRJNA770094, BioSample SAMN30086726. The full code that analyzes the deep mutational scanning experiments is available at https://github.com/jbloomlab/SARS-CoV-2-RBD_MAP_OAS.

## METHODS

### Mice

Wild type C57BL/6J and B6.C(Cg)-*Cd79a*^tm1(cre)Reth^/EhobJ (“*Cd79a*^Cre/+^”, also known as Mb1-Cre[39]) mice were obtained from The Jackson Laboratory. S1pr2-CreERT2 BAC-transgenic mice[40] were a kind gift from T. Kurosaki and T. Okada (U. Osaka, RIKEN-Yokohama). *Igk*^Tag^ mice were generated at the Rockefeller University. We designed the allele as indicated in **Fig. 1a** and **Supplementary Fig. 1**, with a FLAG-tag (DYKDDDDK) and Strep-II tag (WSHPQFEK) separated by stop codons and the SV40 poly-A transcriptional terminator. The 522 nucleotide single-stranded DNA template (including 5’ and 3’ homology arms, each 100 nucleotides long) and the CRISPR guide-RNA (“GGAGCTGGTGGTGGCGTCTC”) were purchased from IDT and prepared according to the Easi-CRISPR gene targeting method[41] by the Rockefeller University Gene Targeting Resource Center and injected into zygotes obtained from C57BL/6 mice by the Rockefeller University Transgenic Services core facility. We verified that the allele was correctly inserted by Sanger sequencing across the entire locus using genomic primers located outside of the homology arms. To decrease the risk of potential CRISPR off-targets, one founder mouse was back-crossed for at least 5 generations onto C57BL/6J mice prior to use in experiments. To generate *Igk*^FLAG/Strep^ mice, germline-excised (Cre-negative mice displaying Strep^+^ B cell surface staining) mice, obtained from occasional spontaneous germline recombination in *Cd79a*^Cre/+^ *Igk*^Tag^ breedings, were crossed to the parental *Igk*^Tag^ strain. All mice were held at the immunocore clean facility at the Rockefeller University under specific pathogen-free conditions. All mouse procedures were approved by the Rockefeller University’s Institutional Animal Care and Use Committee.

### Immunizations, infections, and treatments

Immune responses were induced in 7- to 12-week-old mice by either: subcutaneous footpad immunization with 10 µg TNP_17_ -KLH (Biosearch, #T-5060) supplemented with 1/3 volume of Imject™ alum (ThermoScientific) or i.p. immunization with 50 µg TNP-KLH prepared with 1/3 volume of either alum or aluminum hydroxide gel (alhydrogel, Invivogen); 20 µg recombinantly produced trimer-stabilized HA (see below) in alhydrogel; intramuscular immunization of quadricep muscles with 3 µg WH1[42] or BA.1 spike mRNA encapsulated in lipid nanoparticles (mRNA-LNP), generated as described below; or by intranasal infection with mouse-adapted PR8 influenza virus produced in embryonated chicken eggs (∼33 PFU, kindly provided by M. Carroll, Harvard University Medical School). In S1pr2-*Igk*^Tag^ mice, the primary immune response was fate-mapped by oral gavage of 200 µl tamoxifen (Sigma) dissolved in corn oil at 50 mg/ml, on days 4 and 8 for the day 12 flow cytometry experiment (**Fig. 1**), on days 4, 8 and 12 for all other immunization experiments, and on days 4, 8, 12, and 16 for the influenza infection experiment (**Fig. 3**). In TNP recall experiments (**Fig. 2c**), boosting was performed identically as the primary immunization, on the indicated time points. For homologous mRNA-LNP experiments (**Fig. 2e**), boosting was performed in the same way as priming, except that some mice the left quadricep muscle was boosted (contralaterally, stratified in **Supplementary Fig. 3d**). For heterologous mRNA-LNP recall experiments (**Fig. 4**) boosting was contralateral in all cases. For HA immunization experiments (**Fig. 3**), homologous or heterologous HA protein boosting was performed ipsilaterally, exactly as the primary immunization. For infection experiments, mice were boosted after 3 months by subcutaneous immunization of the footpad with 5 µg HA_PR8_ or HA_FM1_ prepared with 1/3 volume alhydrogel, as described previously[19]. In all cases, additional booster immunizations were performed identically to the first boost. Blood samples were collected via cheek puncture into microtubes prepared with clotting activator serum gel (Sarstedt, #41.1378.005).

### Generation of recombinant and hapten-conjugated proteins

Recombinant HAs used for immunizations and ELISAs were produced in-house using a CHO cell protein expression system, as described previouosly[19]. Cysteine residues were introduced into the HA sequence to create trimer-stabilizing disulfide bonds, as originally described for H1/A/California/07/2009[43]. We described the production of HA_PR8_ and HA_Ca’09_ before[19]. For HA_FM1_ and HA_NC’99_, the same procedure was followed, including the introduction of trimer-stabilizing mutations. For immunizations, C-terminal domains not native to HA (foldon, Avi-tag, His-tag) were removed by thrombin cleavage and HAs were subsequently FPLC-purified prior to storage in phosphate-buffered saline (PBS). For ELISA, non-thrombin treated FPLC-purified proteins were used. A high affinity IgY-specific mAb obtained from CGG-immunized mice (clone 2.1[44]) was modified for use as a standard for FLAG/Strep ELISA detection. Heavy and light chain constant regions in the original human mAb plasmids[45] were replaced with mouse IgG_1_ and Igκ constant regions and the C-terminus of C_κ_ was modified to encode a LoxP site and Ser-Gly-Gly linker followed by either a FLAG or Strep-tag, yielding C_κ_ chains identical to those produced by *Igk*^Tag^ mice prior to and after recombination, respectively. The mAb-FLAG or mAb-Strep light chain plasmids were transfected together with the heavy chain plasmid into 293F cells and purified using protein-G affinity chromatography as described[44]. To compare affinity maturation between FLAG^+^ and Strep^+^ anti-TNP antibody titers (**Fig. 1h**), custom low and high hapten-Bovine serum Albumin (BSA) conjugations were made in-house. BSA (in PBS; ThermoScientific #77110) at 2.5 mg/ml was incubated with TNP-ε-Aminocaproyl-OSu (Biosearch Technologies #T-1030) in PBS with 20% dimethyl sulfoxide at a molar ratio of either 1:2 or 1:20 for 2 hours at room temperature while rotating. Unconjugated TNP-ε-Aminocaproyl-OSu was removed by dialysis in PBS. Final TNP: BSA conjugation ratios were estimated to be ∼1:1 and ∼1:13 by measuring absorbance at 280 and 348 nm, these reagents are referred to as TNP_1_ -BSA and TNP_13_ - BSA. The BSA concentration was corrected by determining the extinction coefficient for TNP-ε-Aminocaproyl-OSu at 280 nm. Besides in **Fig. 1h**, commercial TNP_4_ -BSA (Biosearch Technologies #T-5050) was used for all other TNP ELISAs, see description below.

### Production of mRNA-LNP

The WH1 S mRNA vaccine was designed based on the SARS-CoV-2 S protein sequence (Wuhan-Hu-1, GenBank: MN908947.3) where the lysine (K) and valine (V) amino acids in positions 986-987 are modified to prolines (P) to obtain a prefusion-stabilized mRNA-encoded immunogen. The BA.1 S amino acid sequence was obtained from the WH1 S by introducing BA.1-specific modifications. Coding sequences of the WH1 and BA.1 S were codon-optimized, synthesized and cloned into an mRNA production plasmid (GenScript) as described[46]. mRNA production and LNP encapsulation was performed as described[46]. Briefly, mRNAs were transcribed to contain 101 nucleotide-long poly(A) tails. m1Ψ-5’-triphosphate (TriLink) instead of UTP was used to generate modified nucleoside-containing mRNAs. Capping of the *in vitro* transcribed mRNAs was performed co-transcriptionally using the trinucleotide cap1 analog, CleanCap (TriLink). mRNA was purified by cellulose (Sigma) purification, as described [47]. All mRNAs were analyzed by agarose gel electrophoresis and were stored frozen at -20°C. Cellulose-purified m1Ψ-containing RNAs were encapsulated in LNP using a self-assembly process as previously described wherein an ethanolic lipid mixture of ionizable cationic lipid, phosphatidylcholine, cholesterol and polyethylene glycol-lipid was rapidly mixed with an aqueous solution containing mRNA at acidic pH[48]. The ionizable cationic lipid and LNP composition are described in the patent application WO 2017/004143. The RNA-loaded particles were characterized and subsequently stored at -80°C at a concentration of 1 μg μl^-1^. The mean hydrodynamic diameter of these mRNA-LNP was ∼80 nm with a polydispersity index of 0.02-0.06 and an encapsulation efficiency of ∼95%.

### Flow cytometry

For flow cytometry of peripheral B cells, blood was collected in microtubes with EDTA to prevent coagulation and treated with ACK buffer (Lonza) to lyse red-blood cells. For lymph node samples, cell suspensions were obtained by mechanical disassociation with disposable micropestles (Axygen). Spleens were homogenized by filtering through a 70-μm cell strainer and treated with ACK buffer. Bone-marrow cells were extracted by centrifugation of punctured tibiae and femurs at up to 10,000 xG for 10 s, then treated with ACK buffer. Cells from each tissue were resuspended in PBS supplemented with 0.5% BSA and 1 mM EDTA and incubated first with FC-block (rat anti-mouse CD16/32, clone 2.4G2, Bio X Cell) for 30 min on ice and subsequently with various fluorescently-labeled antibodies (see **Table S1**) for 30 min. Cells were filtered and washed with the same buffer before analysis on a BD FACS Symphony cytometer. Data were analyzed using FlowJo v.10 software.

### Western blotting

To determine the presence of epitope-tagged antibodies in *Igk*^Tag^ mice, serum samples from steady state adult mice and precision plus dual color protein standards (Biorad) were run in triplicate on SDS-PAGE mini-protean TGX protein gels (Biorad) under denaturing conditions, to separate heavy and light antibody chains. Samples were transferred to a PVDF membrane using the Iblot gel transfer system (Invitrogen). Membranes were blocked for 2 hours at room temperature while gently shaking with 5% nonfat dry milk in PBS-Tween (0.05%), prior to overnight incubation in the same buffer with 1:2000 anti-FLAG-HRP (clone D6W5B, CellSignallingTechnology #86861S) or anti-Strep (clone Strep-tag II StrepMAB-Classic, Biorad #MCA2489P) or goat anti-mouse Igκ-HRP (SouthernBiotech #1050-05). Membranes were extensively washed with PBS-Tween and subsequently incubated with western blotting ECL substrate (Amersham) prior to chemiluminescence detection on an Azure c300 gel imager (Azure Biosystems).

### ELISA

ELISAs were performed as described[19], with specific modifications to allow for direct FLAG/Strep comparison. FLAG/Strep ELISAs were performed side by side and with internal standards on each 96-well plate. To detect antigen-specific serum antibody titers, plates were coated overnight at 4°C with antigen in PBS (10 µg/ml for TNP_4_ -BSA, 2 µg/ml for in-house conjugated TNP_1/13_-BSA (see above), 1 µg/ml for HAs and SARS-CoV-2 Spike or RBD proteins (Sinobiological #40592-V08H, *#*40592-V08H121, #40589-V08H26; WH1 S and RBD proteins were a kind gift from P. Wilson)). For FLAG/Strep standard curves, wells were coated with 10 µg/ml purified IgY (Gallus Immunotech). After washing with PBS-Tween (PBS + 0.05% Tween20, Sigma), plates were blocked for 2 hours at room temperature with 2.5% BSA in PBS. Serum samples were diluted 1:100 in PBS and serially titrated in 3-fold dilutions. Mouse anti-IgY mAb-FLAG or mAb-Strep were also serially titrated in 3-fold dilutions (**Supplementary Fig. 2d**). Samples were incubated for 2 hours and then washed with PBS-Tween, before adding one of the following HRP-detection Abs: goat anti-mouse IgG (Jackson Immunoresearch #15-035-071), rat anti-mouse Igκ (abcam #ab99632), goat anti-mouse IgM (Southern Biotech #1020-05), rabbit anti-FLAG-HRP (clone D6W5B) or mouse anti-Strep (clone Strep-tag II StrepMAB-Classic) for 30-45 minutes. Dilutions of anti-FLAG and anti-Strep antibodies were defined so that the curves generated by titration of FLAG- and Strep-tagged mAbs were equivalent (**Supplementary Fig. 2d**). After washing with PBS-Tween, samples were incubated with 3,3’,5,5’-Tetramethylbenzidine substrate (slow kinetic form, Sigma) and the reaction was stopped with 1N HCl. Optical Density (OD) absorbance was measured at 450 nm on a Fisher Scientific accuSkan FC plate reader. To normalize FLAG and Strep endpoint titers, the serum titer dilution was calculated at which each sample passed the threshold OD value of its respective mAb at a fixed concentration of either 20 or 6.67 ng/µl. Titers were calculated by logarithmic interpolation of the dilutions with readings immediately above and immediately below the mAb OD used[49].

For total serum IgG ELISAs, plates were coated with anti-mouse IgG. Standard curves were generated using unlabeled mouse IgG (Southern Biotech #0107-01), and detection was performed with anti-mouse IgG-HRP (Southern Biotech). To deplete IgM from serum samples, anti-mouse IgM agarose beads (Sigma #A4540) were used according to instructions from the manufacturer. Beads were washed with PBS and samples were incubated at a ratio of 1:20 sample to beads overnight at 4°C with rotation. The bead-bound IgM fraction was removed by centrifugation for 3 minutes at 10,000 G, and the unbound supernatant fraction was used for subsequent ELISAs. To confirm the efficiency of IgM depletion, total IgM levels were measured as described above for total IgG, with goat anti-mouse IgM, unlabeled IgM and anti-mouse IgM-HRP (Southern Biotech).

### Serum fractionation and SARS-Cov2 pseudoneutralization assay

Virus neutralization titers were assessed in FLAG^+^ versus Strep^+^ serum fractions of samples collected from S mRNA-LNP immunized S1pr2-*Igk*^Tag^ mice. To separate fractions, immunoprecipitation with anti-FLAG M2 magnetic beads (Sigma #M8823-5ML) and MagStrep “type3” XT beads (IBA #2-4090-010) was performed as per the manufacturer’s instructions. In brief, magnetic beads were washed with sample buffer (Tris buffered saline for FLAG beads, and 1x buffer W (IBA # 2-1003-100) for MagStrep beads) and samples were incubated at a ratio of 20:1 sample to bead resin overnight at 4°C with rotation. Bead-bound fractions were separated using a magnetic separator and discarded, while the unbound fraction was collected. Fractionated samples were concentrated by centrifugation to half the input concentration and heat inactivated. The degree to which the total and fractionated serum samples neutralized WH1 and BA.1 SARS-CoV-2 was approximated using SARS-CoV-2 spike pseudotyped HIV-1 based NanoLuc luciferase reporter assay described previously[24]. Briefly, serum samples were five-fold serially diluted with a final top dilution of 1:00 serum and incubated for 1 h at 37°C with SARS-CoV-2 WH1 or BA.1 spike pseudotyped HIV-1 reporter virus and then transferred to HT1080/ACE2.cl14 cells[50]. At 48 h, the cells were washed, lysed and luciferase activity was measured using the Nano-Glo Luciferase Assay System (Promega) and the Glomax Navigator luminometer (Promega). The relative luminescence units were normalized using cells infected in the absence of serum and then plotted in GraphPad Prism. NT50 values were calculated using four-parameter non-linear regression (least squares regression method without weighting) of the curves shown in **Supplementary Fig. 5b**. The mean of two technical duplicates is shown, outlier points were excluded. For NT50 comparisons between input and fractions (**Fig. 4g** and **Supplementary Fig. 5c**), the NT50 of the fractionated samples was adjusted to equalize the BA.1 RBD ELISA titer of the un-depleted tag compared to its corresponding ELISA titer in the input fraction.

### Deep Mutational Scanning

#### Construction of yeast-displayed deep mutational scanning libraries of Omicron BA.1 RBD

Duplicate single-mutant site-saturation variant libraries were designed in the background of the SARS-CoV-2 Omicron BA.1 spike RBD and produced by Twist Bioscience, essentially the same as has been done previously for other SARS-CoV-2 variants[51, 52]. The Genbank map of the plasmid encoding the unmutated Omicron BA.1 RBD in the yeast-display vector is available at https://github.com/jbloomlab/SARS-CoV-2-RBD_DMS_Omicron/blob/main/data/3294_pETcon-SARS2-RBD_Omicron-BA1.gb. The site-saturation variant libraries were delivered as double-stranded DNA fragments by Twist Bioscience and were barcoded and cloned in bulk into the yeast-display vector backbone. The barcoded mutant library plasmid DNA was electroporated into *E. coli* (NEB 10-beta electrocompetent cells, New England BioLabs #C3020K), and bottlenecked to ∼1 × 10^5^ cfus (an average of >25 barcodes per single-mutant). Plasmid DNA was purified and transformed into the AWY101 yeast strain. 16-nucleotide barcodes were associated with their BA.1 variants by PacBio sequencing, and the effects of mutations of RBD expression and ACE2 binding were measured, essentially as described[52]. These experiments are described and analyzed at https://github.com/jbloomlab/SARS-CoV-2-RBD_DMS_Omicron.

#### FACS sorting of yeast libraries to select mutations with reduced binding by polyclonal sera from immunized mice

Experiments mapping mutations that reduce RBD binding of sera from immunized mice were performed in biological duplicate with independent mutant WH1 or BA.1 RBD libraries, similarly to as previously described for monoclonal antibodies[53], human polyclonal plasma samples[54]. First, 75 µL of each of the sera was twice-depleted of nonspecific yeast-binding antibodies by incubating for 2 hours at room temperature or overnight at 4°C with 37.5 OD units of AWY101 yeast containing an empty vector, as described[54]. WH1 and BA.1 mutant RBD yeast libraries[52] were induced with galactose-containing, low-dextrose synthetic defined medium with casamino acids (SD-CAA, 6.7g/L Yeast Nitrogen Base, 5.0g/L Casamino acids, 1.065 g/L MES acid, and 2% w/v galactose + 0.1% w/v dextrose) to express RBD, then washed and incubated with diluted serum for 1 hour at room temperature with gentle agitation. Each tested combination of mouse serum against each WH1 or BA.1 RBD mutant library for loss of binding of Strep or FLAG-tag antibodies was performed independently. For each serum, a sub-saturating dilution was used such that the amount of fluorescent signal due to serum antibody binding to RBD was approximately equal across samples (1:1000 for mapping of Strep antibodies against the WH1 libraries, 1:200 for mapping of Strep antibodies against the BA.1 libraries, and 1:50 for mapping of the FLAG antibodies against the BA.1 libraries). The yeast libraries were then secondarily labeled for 1 hour with 1:100 FITC-conjugated anti-MYC antibody (Immunology Consultants Lab, #CYMC-45F) to label for RBD expression and either 1:200 APC-conjugated Streptavidin (Invitrogen S-868) to label for bound Strep antibodies or APC-conjugated rat anti-FLAG (BioLegend #637308) to label for bound FLAG-tagged antibodies. A flow cytometric selection gate was drawn to capture RBD mutants with reduced antibody binding for their degree of RBD expression. For each sample, ∼4 × 10^6^ cells were processed on the BD FACSAria II cell sorter. Serum-escaped cells were grown overnight in SD-CAA as defined above with 2% w/v dextrose, no galactose, and 100 U/mL penicillin + 100 µg/mL streptomycin to expand cells prior to plasmid extraction.

#### DNA extraction and Illumina sequencing

Plasmid DNA was extracted from 30 OD units (1.6 × 10^8^ colony forming units (cfus)) of pre-selection yeast populations and approximately 5 OD units (∼3.2 × 10^7^ cfus) of overnight cultures of serum-escaped cells (Zymoprep Yeast Plasmid Miniprep II) as previously described[51, 53]. The 16-nucleotide barcodes identifying each WH1 or BA.1 RBD variant were amplified by polymerase chain reaction (PCR) and prepared for Illumina sequencing as described previously[51, 53]. Barcodes were sequenced on an Illumina NextSeq 2000 with 50 bp single-end reads.

#### Analysis of deep sequencing data to compute each mutation’s escape fraction

Escape fractions were computed essentially as described[53]. We used the dms_variants package (https://jbloomlab.github.io/dms_variants/, version 1.4.0) to count each barcoded RBD variant in each pre-selection and serum-escape population. For each selection, we computed the escape fraction for each barcoded variant via the formula provided in Greaney et al.[53]. These escape fractions represent the estimated fraction of cells expressing that specific variant that falls in the escape bin, such that a value of 0 means the variant is always bound by serum and a value of 1 means that it always escapes serum binding. We then applied a computational filter to remove variants with >1 amino-acid mutation, low sequencing counts (< 50 in the pre-selection condition), or highly deleterious mutations that might cause antibody escape simply by leading to poor expression of properly folded RBD on the yeast cell surface (an ACE2 binding score of < –2 or an RBD expression score of < –1.25 or –0.83361 for the WH1 and BA.1 mutant libraries, respectively, reflecting the different baseline expression levels of the two wild-type RBDs). The reported antibody-escape scores throughout the paper are the average across duplicate libraries; these scores are also in **Supplementary Spreadsheet 1**. Correlations in final single-mutant escape scores are shown in Supplementary Fig 6c. Full documentation of the computational analysis is at https://github.com/jbloomlab/SARS-CoV-2-RBD_MAP_OAS.

#### Data visualization

The serum-escape map logo and line plots were created using the dmslogo package (https://jbloomlab.github.io/dmslogo, version 0.6.2). The height of each letter indicates the escape fraction for that amino-acid mutation. For each serum, the logo plots feature any site where for >=1 library/antibody tag condition, the site-total antibody escape was >10x the median across all sites and at least 10% the maximum of any site. For each sample, the y-axis is scaled to be the greatest of (a) the maximum site-wise escape metric observed for that sample, or (b) 20x the median site-wise escape fraction observed across all sites for that plasma. The code that generates these logo plot visualizations is available at https://github.com/jbloomlab/SARS-CoV-2-RBD_MAP_OAS/blob/main/results/summary/escape_profiles.md. To visualize serum escape on the RBD structure, the WH1 RBD surface (PDB 6M0J) was colored by the site-wise escape metric at each site, with white indicating no escape and red indicating the site with the most escape.

### Statistical analysis and software

Statistical tests used to compare conditions are indicated in figure legends. Statistical analysis was carried out using GrahPad Prism v.9. Flow cytometry analysis was carried out using FlowJo v.10 software. Graphs were plotted using Prism v.9, and edited for appearance using Adobe Illustrator CS. For data plotted on logarithmic scales (e.g., serum antibody titers), statistical analysis was performed on the log-transformed data. Samples with reactivities below the limit of detection were assigned a value of 100, as the top dilution was 1:100.

## SUPPLEMENTARY FIGURES AND LEGENDS

**Supplementary Figure 1.**
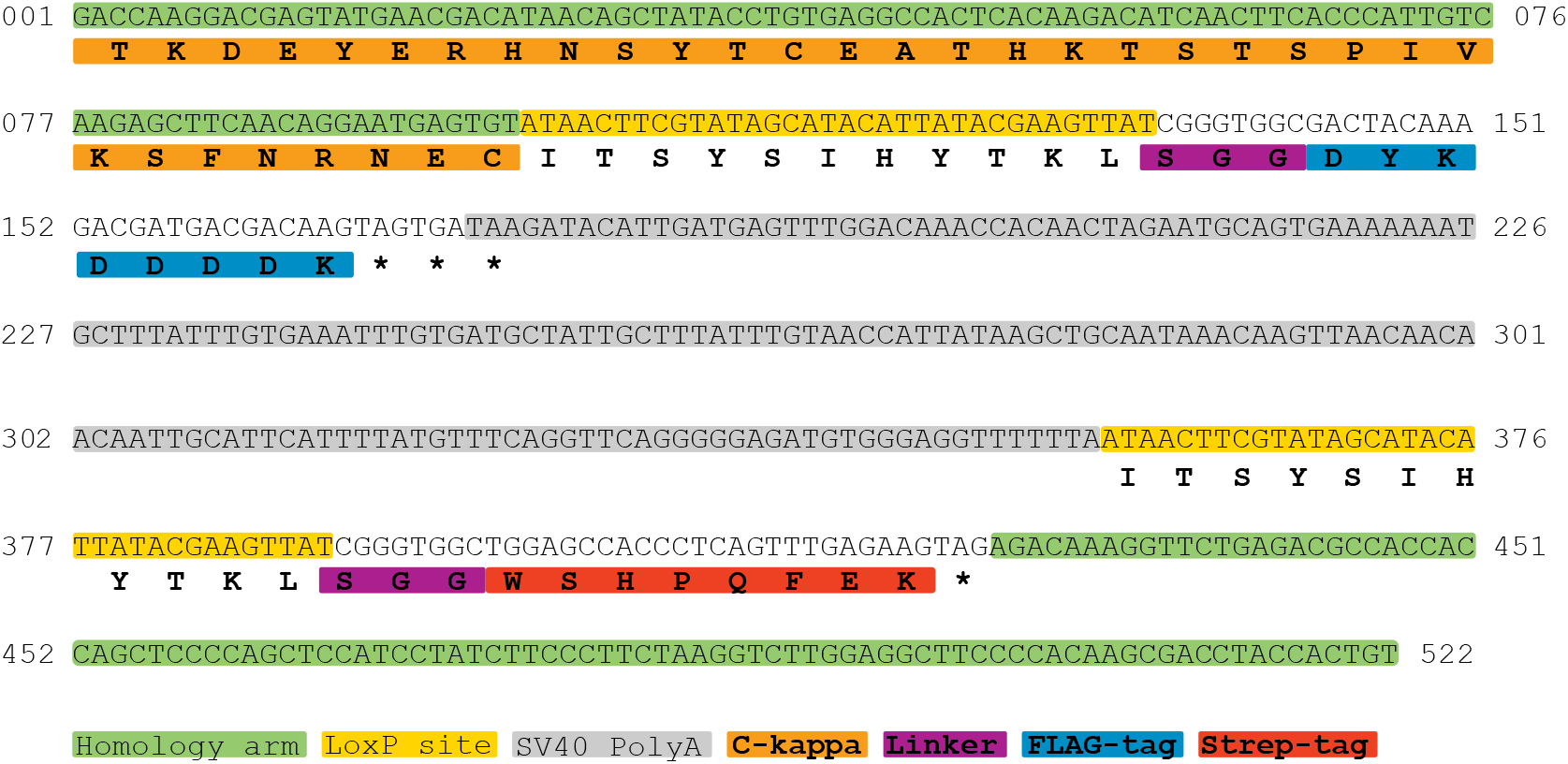
Design of the Κ-tag allele. Nucleotide sequence of the 522 bp DNA template used to generate the *Igk*^Tag^ allele, with aa translations given for all coding sequences (bold font). Nucleotide numbers are given for each line. Amino acid translation is positioned below the center nucleotide of each codon.

**Supplementary Figure 2.**
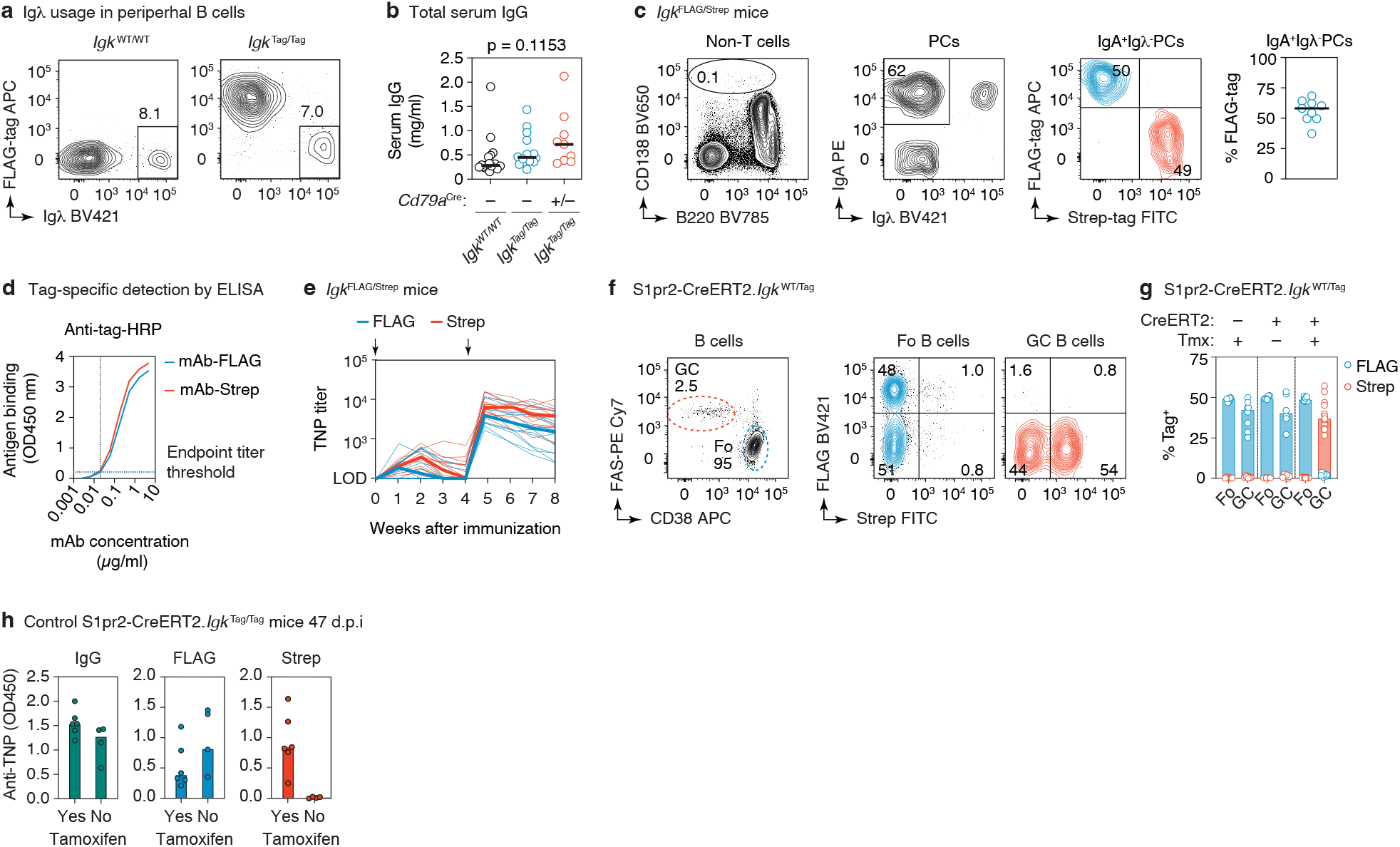
Characterization of the Κ-tag system. **(a)** Representative flow cytometry plots of peripheral B cells (gated: B220^+^CD4^−^CD8^−^CD138^−^) obtained from the blood of WT and *Igk*^Tag/Tag^ mice, stained for surface expression of Igλ and FLAG-tagged immunoglobulins. **(b)** Total IgG concentration in the serum of WT, *Igk*^Tag/Tag^, and *Cd79a*^Cre/+^.*Igk*^Tag/Tag^ mice as determined by ELISA. Differences were not significant by one-way ANOVA (p<0.05). **(c)** Flow cytometry of steady-state bone marrow plasma cells (PCs) obtained from adult (6 week old) *Igk*^FLAG/Strep^ mice. Gating strategy for PCs is shown in the left panel (pre-gated on non-CD4/CD8 T cells), for surface IgA^+^ and Igλ^−^ PCs in the middle panel, and for FLAG/Strep in the right panel. Quantification across multiple mice is shown in the rightmost panel (each dot represents an individual *Igk*^FLAG/Strep^ mouse, line represents the median). **(d)** ELISA standard curves with monoclonal antibody (mAb)-FLAG and mAb-Strep detected at dilutions of the respective HRP antibodies where the curves overlap. The mAb concentration at which the curves crossed the absorbance background threshold (indicated by the dotted lines) was used to calculate the endpoint titer, as described in the methods section. **(e)** Tag-specific anti-TNP titers in *Igk*^FLAG/Strep^ mice immunized and boosted i.p. with TNP-KLH/alum at the timepoints indicated by black arrows. Results are from 14 mice from 2 independent experiments. The day 7 timepoint was not collected for the first cohort. Thin lines represent individual mice, thick lines link the medians of the log transformed titer values at each time point. **(f)** Flow cytometry of S1pr2-*Igk*^WT/Tag^ mice as in Fig. 1d, e. with quantification in **(g)**, cre^−^ and no tamoxifen control groups are included. **(h)** Anti-TNP ELISA reactivity for S1pr2-*Igk*^Tag/Tag^ mice immunized as in Fig. 1d. Serum was obtained at 47 d.p.i from mice that received tamoxifen (same mice as Fig. 1g) and that did not receive tamoxifen. IgG (left), FLAG (middle) and Strep (right) ELISA absorbance is shown. Samples were diluted 1:100.

**Supplementary Figure 3.**
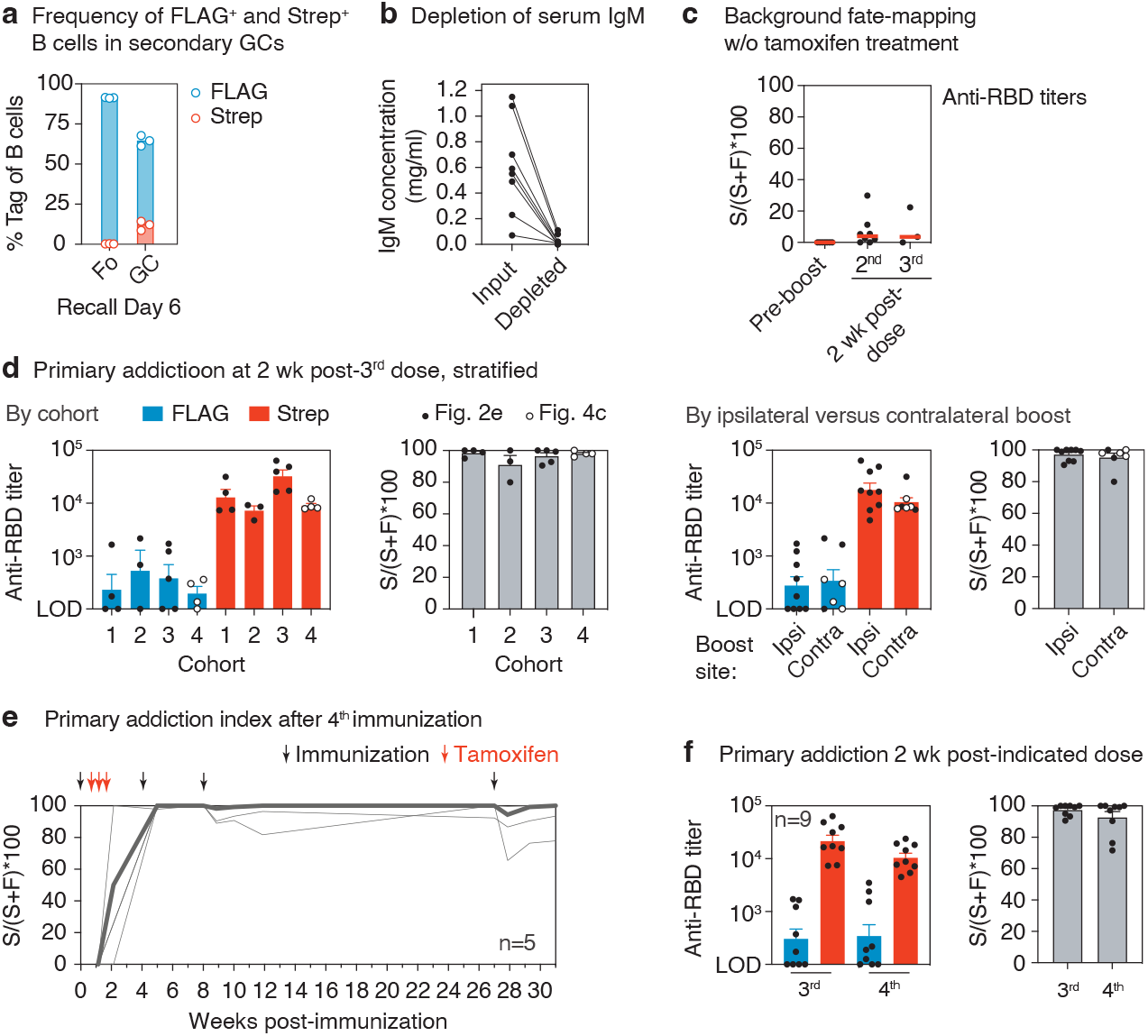
Recall antibodies and GCs. **(a)** Flow cytometry of secondary GCs in the spleen of TNP-KLH i.p. primed and boosted S1pr2-*Igk*^Tag/Tag^ mice, with tamoxifen labeling at 4, 8, and 12 d.p.i.. Gated on GC B cells (FAS^+^CD38^−^B220^+^CD4^−^CD8^−^CD138^−^) expressing FLAG or Strep-tag as in Fig. 1e. **(b)** Total IgM concentrations of serum samples pre and post IgM-depletion via immunoprecipitation, measured by ELISA. **(c)** Background Strep^+^ anti-RBD titers in S1pr2-*Igk*^Tag/Tag^ control mice immunized as in **Fig. 2a,e**, but not treated with tamoxifen. Graphs show median percentage of the anti-RBD titer that is Strep^+^ ((S/(S+F)*100)) in the absence of tamoxifen at the pre-boost time point and two weeks after the second and third immunizations. This represents the median percentage by which FLAG^+^ titers in recall responses are likely to be underestimated by spontaneous recombination by the S1pr2-CreERT2 driver. **(d)** Comparison of primary imprinting data shown in Fig. 2e,f, stratified by cohort and ipsilateral versus contralateral boost. The 4^th^ cohort of mice is that the homologously boosted group showin in Fig. 4b-e, depicted here by open circles. Bars represent the mean of log transformed titers, error bars are SEM. **(e)** Primary addiction index after the 4^th^ immunization with SARS-CoV-2 S protein mRNA-LNP, based on ELISA data shown in Fig. 2g. **(f)** Comparison of primary addiction between 3^rd^ and 4^th^ responses, based on data from (e) and Fig. 2e,f.

**Supplementary Figure 4.**
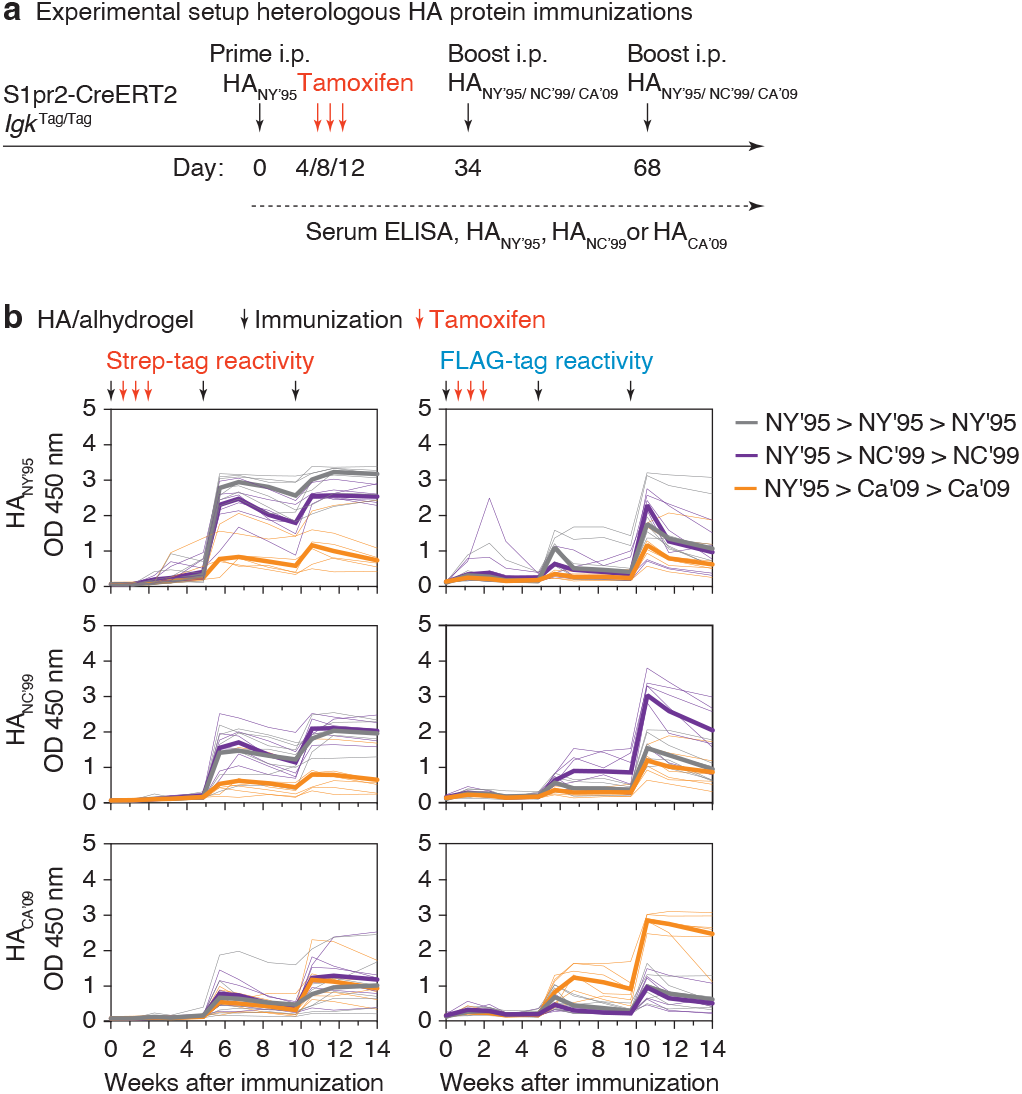
Primary cohort recall upon heterologous HA protein boosting. **(a)** Schematic representation of HA prime-boost strategy. S1pr2-*Igk*^Tag/Tag^ mice were primed i.p. with HA_NY’95_ in alhydrogel and boosted homologously or heterologously with HA_NC’99_ or HA_CA’09_ in alhydrogel as indicated. **(b)** Full time course of anti-HA tag-specific ELISA reactivity (optical density at 1:100 dilution) of the same mice shown in Fig. 3f. Anti-HA_NY’95_ (top), HA_NC’99_ (middle) and HA_CA’09_ (bottom) ELISAs are shown, with Strep (left) and FLAG (right) detection.

**Supplementary Figure 5.**
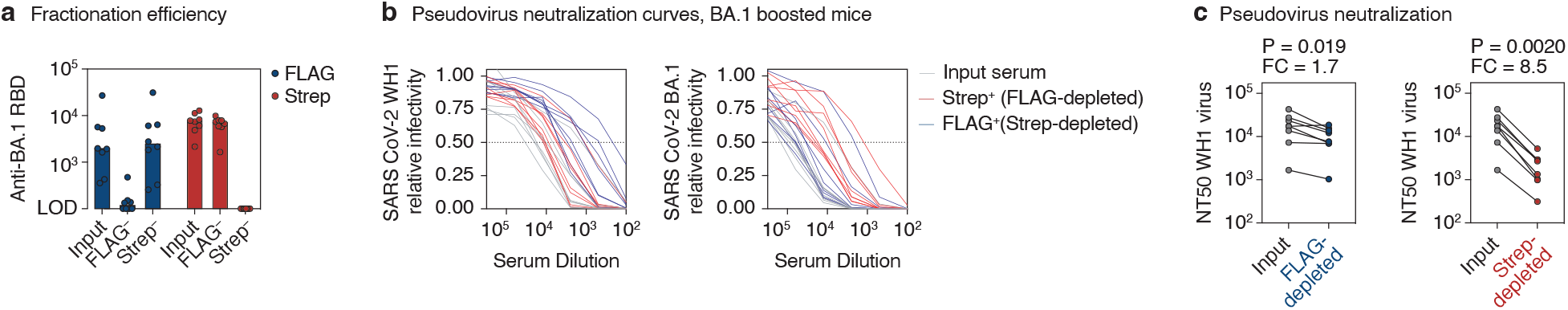
Neutralization of WH1 and BA.1 pseudoviruses by serum antibody fractions from BA.1 boosted mice. **(a)** Efficiency of serum fractionation into FLAG and Strep-depleted fractions, as measured by anti-RBD ELISA of input vs. post-depletion samples. Serum was obtained from heterologously immunized mice two weeks after the 3^rd^ immunization, Fig. 4d. **(b)** Neutralization of WH1 (left) and BA.1 (right) SARS-CoV-2 S-expressing pseudotyped HIV-1 virus by serum fractions shown in (a). Mean values of technical duplicates are shown. **(c)** WH1 S pseudovirus NT50 titers for samples in (b). Post-depletion NT50s were normalized to input serum based on the BA.1 RBD ELISA titers (a), by applying a correction equalizing the anti-RBD Strep-titer of the FLAG-depleted fraction to the input, and the anti-RBD FLAG-titer of the Strep-depleted fraction to the input. P-values are for one-tailed paired T test. FC, fold-change.

**Supplementary Figure 6.**
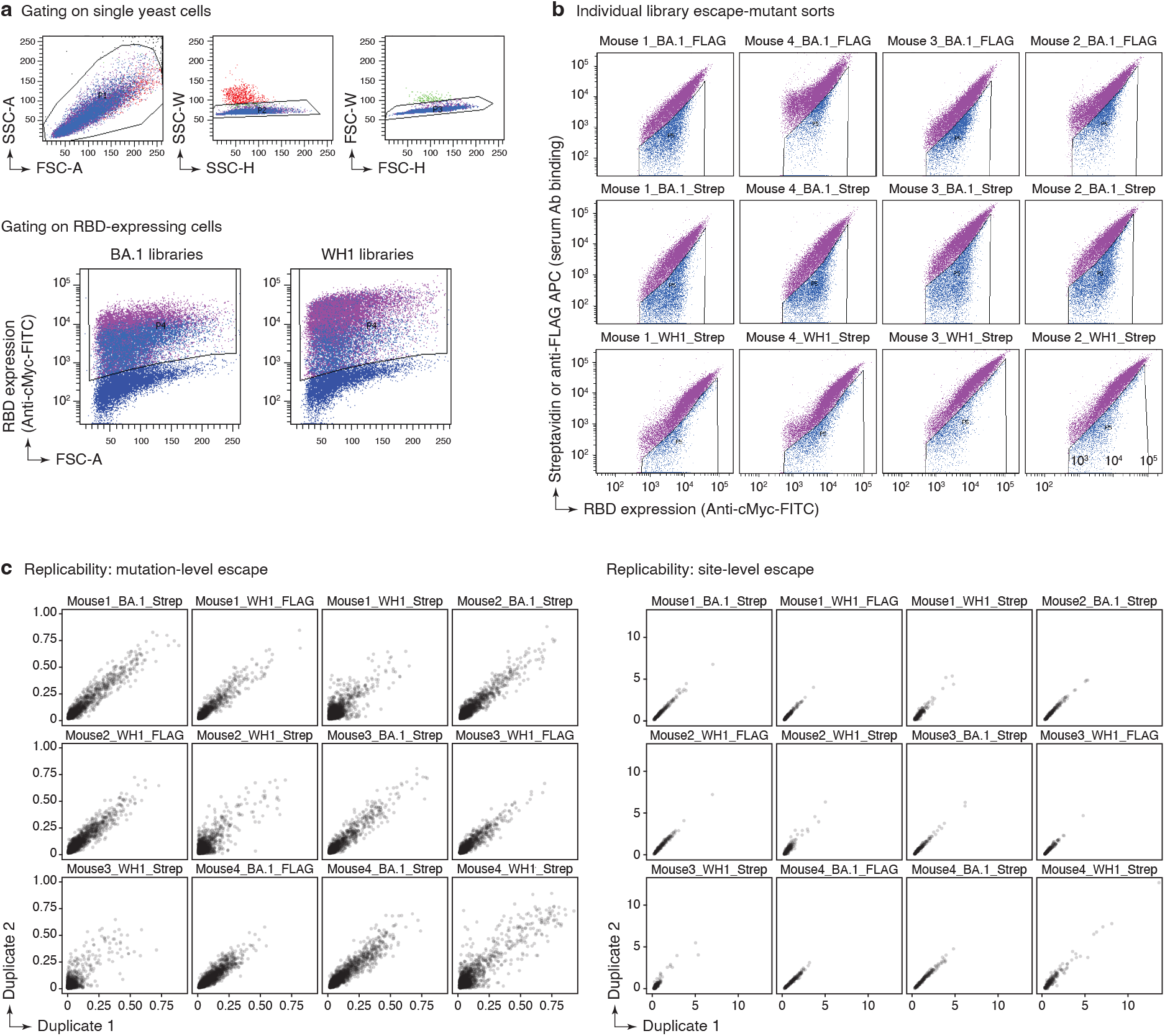
Yeast-displayed deep mutational scanning to map mutations that reduce binding of immunized mouse serum. **(a)** Top: Representative plots of nested FACS gating strategy used for all experiments to select for single yeast cells. Bottom: Gating strategy to select for RBD-expressing single cells (FITC-A vs. FSC-A). **(b)** FACS gating strategy for one of two independent libraries to select cells expressing BA.1 or WH1 RBD mutants with reduced Strep or FLAG antibody binding (cells in blue), as measured by secondary staining with APC-conjugated streptavidin or APC-conjugated anti-FLAG antibody, respectively. Gates were set manually for each sample to capture cells that have a reduced amount of tagged antibody binding for their degree of RBD expression. FACS scatter plots were qualitatively similar between the two libraries. The mouse identifier (#1-4), DMS target library (WH1 or BA.1), and antibody tag (Strep or FLAG) are indicated above each plot. **(c)** Mutation (left)- and site (right)-level correlations of escape scores between two independent biological replicate libraries.

**Supplementary Figure 7.**
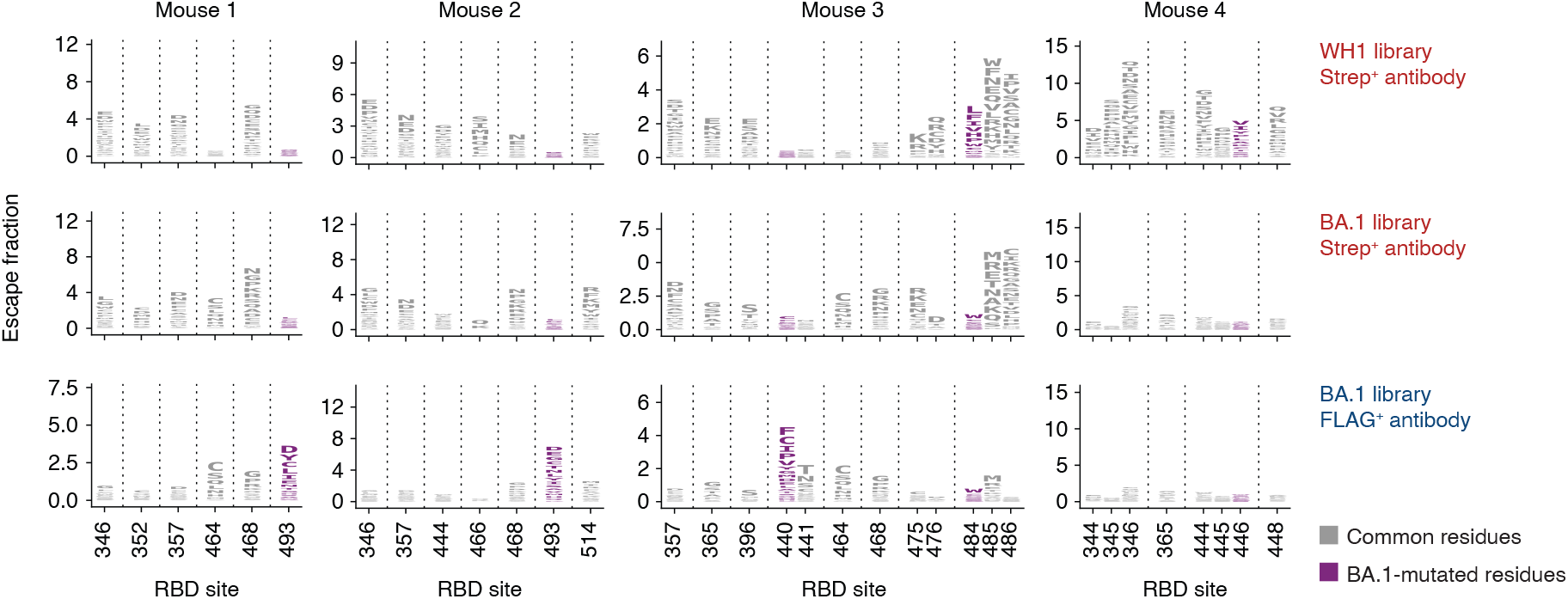
RBD serum-escape logo plots. Deep mutational scanning results of serum collected from 4 heterologously immunized mice, 2 weeks after the 3^rd^ dose (Fig. 4d). Each mutation’s “escape fraction” was measured, which ranges from 0 (no cells effect on antibody binding) to 1 (all cells with the mutation have decreased antibody binding). Mouse 4 is not shown in the main text as there were no interpretable peaks in antibody binding to BA.1 libraries for either antibody fraction. Logo plots show the antibody-escape fractions for individual amino-acid mutations at key sites of strong escape. Sites in which BA.1 differs from the WH1 sequence are shown in purple font. All escape scores are shown in **Supplementary Spreadsheet 1** and are available online at https://github.com/jbloomlab/SARS-CoV-2-RBD_MAP_OAS/blob/main/results/supp_data/all_raw_data.csv.

**Table S1.** Antibodies used for flow cytometry.

| **Antibody** | **Clone** | **Fluorophore** | **Final Dilution** | **Company** | **Catalog number** |
| --- | --- | --- | --- | --- | --- |
| B220 | RA3-6B2 | BV785 | 1:400 | Biolegend | 103246 |
| CD4 | RM4-5 | V500 | 1:200 | BD Biosciences | 560782 |
| CD8a | 53-6.7 | V500 | 1:200 | BD Biosciences | 560776 |
| CD38 | 17-0381-82 | APC | 1:400 | eBioscience | 17-0381-82 |
| Fas/CD95 | Jo2 | PE/Cy7 | 1:800 | BD Biosciences | 557653 |
| CD138 | 281-2 | BV650 | 1:400 | Biolegend | 142517 |
| FLAG (DYKDDDDK) | L5 | APC | 1:200 | Biolegend | 637308 |
| FLAG (DYKDDDDK) | L5 | BV421 | 1:200 | Biolegend | 637322 |
| Strep | 5A9F9 | FITC | 1:200 | LS Bio | LS-C203631-100 |
| IgLambda | R26-46 | BV421 | 1:200 | BD Biosciences | 744523 |
| IgLambda | R26-46 | FITC | 1:200 | BD Biosciences | 553434 |
| IgA | mA-6E1 | PE | 1:200 | eBioscience | 12-4204-82 |
| IgM | II/41 | APC-eFluor 780 | 1:200 | eBioscience | 47-5790-82 |
| IgM | II/41 | PE/Cy7 | 1:200 | eBioscience | 25-5790-81 |
| CD16/32 | 2.4G2 | NA | 1:200 | Bio X Cell<br>Immunology | BE0307 |
| MYC | Polyclonal | FITC | 1:100 | Consultants Lab | CYMC-45F |
| Streptavidin | NA | APC | 1:200 | Invitrogen | S-868 |

